# A polygenic risk score for breast cancer in U.S. Latinas and Latin-American women

**DOI:** 10.1101/598730

**Authors:** Yiwey Shieh, Laura Fejerman, Paul C. Lott, Katie Marker, Sarah D. Sawyer, Donglei Hu, Scott Huntsman, Javier Torres, Magdalena Echeverry, Mabel E. Bohorquez, Juan Carlos Martínez-Chéquer, Guadalupe Polanco-Echeverry, Ana P. Estrada-Florez, the COLUMBUS Consortium, Christopher A. Haiman, Esther M. John, Lawrence H. Kushi, Gabriela Torres-Mejía, Tatianna Vidaurre, Jeffrey N. Weitzel, Sandro Casavilca Zambrano, Luis G. Carvajal-Carmona, Elad Ziv, Susan L. Neuhausen

## Abstract

**Background:** Over 180 single nucleotide polymorphisms (SNPs) associated with breast cancer susceptibility have been identified; these SNPs can be combined into polygenic risk scores (PRS) to predict breast cancer risk. Since most SNPs were identified in predominantly European populations, little is known about the performance of PRS in non-Europeans. We tested the performance of a 180-SNP PRS in Latinas, a large ethnic group with variable levels of Indigenous American, European, and African ancestry.

**Methods:** We conducted a pooled case-control analysis of U.S. Latinas and Latin-American women (4,658 cases, 7,622 controls). We constructed a 180-SNP PRS consisting of SNPs associated with breast cancer risk (p < 5 × 10^−8^). We evaluated the association between the PRS and breast cancer risk using multivariable logistic regression and assessed discrimination using area under the receiver operating characteristic curve (AUROC). We also assessed PRS performance across quartiles of Indigenous American genetic ancestry.

**Results:** Of 180 SNPs tested, 142 showed directionally consistent associations compared with European populations, and 39 were nominally significant (p < 0.05). The PRS was associated with breast cancer risk, with an odds ratio (OR) per standard deviation increment of 1.58 (95% CI 1.52 to 1.64) and AUCROC of 0.63 (95% CI 0.62 to 0.64). The discrimination of the PRS was similar between the top and bottom quartiles of Indigenous American ancestry.

**Conclusions:** The 180-SNP PRS predicts breast cancer risk in Latinas, with similar performance as reported for Europeans. The performance of the PRS did not vary substantially according to Indigenous American ancestry.

## Introduction

Over 180 single nucleotide polymorphisms (SNPs) associated with breast cancer susceptibility have been discovered in genome-wide association studies (GWAS) [1–4]. Though each SNP has a modest effect, multiple SNPs can be combined into a polygenic risk score (PRS) [5]. PRS has emerged as a promising tool for breast cancer risk stratification. The risk associated with having a PRS in the upper 20-25^th^ percentile is similar to that of strong clinical risk factors such as having extremely dense breasts [6], and adding PRS to risk models improves discrimination and reclassification [6–8]. Ongoing clinical trials are studying the use of PRS to personalize breast cancer screening and prevention [9]. Some commercial genetic testing laboratories are already returning PRS results to those who tested negative for pathogenic moderate- or high-penetrance mutations [10, 11].

A major barrier to the widespread use of PRS is the paucity of knowledge regarding its performance in non-European populations. To date, SNP discovery has overwhelmingly occurred in European populations [12]. However, the effect sizes, allele frequencies, and linkage disequilibrium patterns of SNPs vary by ancestry [12, 13]. Though relatively few studies have examined PRS performance in non-Europeans, they suggest that PRS constructed using European SNP summary statistics (effect size, allele frequency) perform worse in Latinas [14] and women of African ancestry [14, 15]. Currently, commercial testing laboratories only report breast cancer PRS results to women of European ancestry [10, 11].

Disparities in the use and performance of PRS could especially affect Latinas. Latino/Latinas comprise the largest minority group in the U.S., representing 17.8% of the population in 2016 [16]. This group includes genetically admixed individuals who have varying degrees of Indigenous American, European, African, and Asian ancestry [17–19]. We previously identified SNPs in the 6q25 locus associated with breast cancer risk exclusively in Latinas [20]. Most SNPs discovered in European populations display directional consistency in Latinas, with some being nominally significant [20, 21]. One previous study assessed the performance of a breast cancer PRS in Latinas, finding that a 71-SNP PRS had worse prediction in Latinas compared to Europeans [5, 14]. However, it included only 147 cases and did not account for genetic ancestry [14].

We sought to test the performance of PRS in U.S. Latinas and Latin American women (collectively referred to hereafter as *Latinas*). To that end, we conducted a pooled case-control analysis of 8 studies comprising 13,624 Latinas. We examined the predictive performance of a 71-SNP and a 180-SNP PRS, and whether PRS performance varies by genetic ancestry.

## Methods

### Participants

Our analysis included 13,624 self-identified Latinas, of whom 5,697 women with invasive breast cancer were considered cases and 7,927 without breast cancer were controls. Participants came from 8 studies (Tables 1 **and S1**). Recruitment details and patient characteristics have been previously reported for each study except for PEGEN-BC. Studies are briefly described below and in more detail in the **Supplement**.

**Table 1.** Participant characteristics by study and case-control status

|  | **SFBCS/NC-BCFR** |  | **Kaiser RPGEH** |  | **MEC** |  | **CAMA** |  | **COLUMBUS**<br>**(Colombia)** |  | **COLUMBUS**<br>**(Mexico)** |  |
| --- | --- | --- | --- | --- | --- | --- | --- | --- | --- | --- | --- | --- |
|  | Controls | Cases | Controls | Cases | Controls | Cases | Controls | Cases | Controls | Cases | Controls | Cases |
| Number of individuals | 589 | 942 | 3563 | 222 | 1469 | 532 | 702 | 709 | 761 | 954 | 453 | 481 |
| Age at diagnosis (cases) or<br>interview (controls) in<br>years, mean (SD) | 53 (11) | 50 (11) | 55 (13) | 57 (10) | 67 (8) | 66 (8) | 52 (9) | 52 (10) | 64 (10) | 52 (10) | 35 (12) | 57 (13) |
| Positive family history of<br>breast cancer, n (%) | 55 (9)* | 190 (20)* | 211 (6)‡ | 38 (17)‡ | 141 (10)* | 73 (14)* | 27 (4) | 50 (7) | ND | 49 (5)† | 34 (8)‡ | 23 (5)‡ |
| Genetic ancestry, mean %<br>(SD) |  |  |  |  |  |  |  |  |  |  |  |  |
| Indigenous American | 39.1<br>(16.9) | 35.9<br>(17.3) | 29.1<br>(18.1) | 26.9<br>(17.0) | 38.6<br>(14.2) | 36.4<br>(13.5) | 63.9<br>(18.1) | 59.0<br>(18.2) | 43.2<br>(10.8) | 43.3<br>(10.0) | 57.0<br>(14.8) | 57.7<br>(17.7) |
| European | 52.6<br>(17.4) | 56.0<br>(17.8) | 62.4<br>(19.8) | 64.5<br>(19.4) | 54.3<br>(15.2) | 56.2<br>(14.1) | 30.9<br>(16.2) | 35.5<br>(17.1) | 50.0<br>(10.6) | 49.5<br>(9.9) | 36.2<br>(14.0) | 37.3<br>(17.0) |
| African | 6.3 (7.5) | 6.2 (6.8) | 5.9 (8.3) | 5.0 (4.4) | 4.9 (2.8) | 5.2 (3.4) | 3.7 (3.2) | 3.8 (3.0) | 6.2 (5.5) | 6.7 (5.1) | 65.8 (2.5) | 4.2 (3.2) |
| Asian | 2.0 (3.7) | 1.9 (3.2) | 2.6 (6.7) | 3.6 (9.1) | 2.3 (3.2) | 2.1 (4.4) | 1.5 (1.3) | 1.7 (2.0) | 0.6 (0.8) | 0.6 (0.7) | 1.3 (2.5) | 0.8 (0.8) |
| Estrogen receptor status<br>Positive§<br>Negative§<br>Unknown | NA | 593 (72)\|\|<br>230 (28)<br>119 | NA | 161 (85)<br>29 (15)<br>32 | NA | 303 (74)<br>108 (26)<br>121 | NA | 116 (69)<br>52 (31)<br>541 | NA | 354 (67)<br>177 (33)<br>423 | NA | 140 (77)<br>41 (23)<br>300 |
| 71-SNP PRS, mean (SD)¶ | 0.99<br>(0.49) | 1.19<br>(0.57) | 1.01<br>(0.50) | 1.21<br>(0.55) | 1.01<br>(0.47) | 1.18<br>(0.52) | 0.93<br>(0.48) | 1.07<br>(0.49) | 0.99<br>(0.45) | 1.20<br>(0.59) | 0.93<br>(0.45) | 1.08<br>(0.55) |
| 180-SNP PRS, mean<br>(SD)# | 1.00<br>(0.68) | 1.26<br>(0.79) | 0.98<br>(0.61) | 1.27<br>(0.64) | 1.03<br>(0.64) | 1.29<br>(0.72) | 0.99<br>(0.71) | 1.23<br>(0.77) | 1.02<br>(0.61) | 1.32<br>(0.80) | 1.00<br>(0.61) | 1.24<br>(0.79) |

|  | PEGEN-BC/Peru |  | COH/CCGCRN |  | All |  |
| --- | --- | --- | --- | --- | --- | --- |
|  | Controls | Cases | Controls | Cases | Controls | Cases |
| Number of individuals | 85 | 818 | 305 | 1039 | 7927 | 5697 |
| Age at diagnosis or interview in years, mean (SD) | ND | 50 (11) | 52 (11) | 43 (9) | 57 (13) | 52 (12) |
| Positive family history of breast cancer, n (%) | ND | 54 (7) | 26 (9)† | 348 (33)† | 494 (6) | 825 (14) |
| Genetic ancestry, mean % (SD) |  |  |  |  |  |  |
| Indigenous American | 76.3 (15.1) | 76.3 (16.3) | 40.8 (18.8) | 43.2 (19.2) | 38.6 (20.2) | 48.7 (21.6) |
| European | 17.8 (11.2) | 17.0 (11.4) | 48.0 (18.0) | 45.4 (18.1) | 53.8 (20.4) | 44.3 (20.7) |
| African | 3.6 (5.9) | 4.4 (7.8) | 4.4 (3.8) | 4.9 (5.8) | 5.5 (6.5) | 5.2 (5.6) |
| Asian | 2.3 (4.5) | 2.3 (4.7) | 2.4 (2.1) | 2.9 (3.8) | 2.1 (4.9) | 1.9 (3.7) |
| Estrogen receptor status<br>Positive§<br>Negative§<br>Unknown | NA | 548 (69)<br>246 (31)<br>24 | NA | 585 (72)<br>233 (28)<br>221 | NA | 2800 (72)<br>1116 (28)<br>1781 |
| 71-SNP PRS, mean (SD)\|\| | 0.93 (0.46) | 1.16 (0.54) | 0.99 (0.48) | 1.16 (0.55) | 0.99 (0.48) | 1.16 (0.55) |
| 180-SNP PRS, mean (SD)¶ | 1.08 (0.73) | 1.44 (0.95) | NA | NA | 1.00 (0.64) | 1.30 (0.81) |

Abbreviations: CAMA = Cancer de Mama; COH/CCGCRN = Clinical Cancer Genetics Community Research Network; COLOMBUS = Colombian Study of Environmental and Heritable Causes of Breast Cancer; MEC = Multiethnic Cohort; NC-BCFR = Northern California Breast Cancer Family Registry; ND = not determined; PRS = polygenic risk score; RPGEH = Research Project on Genes, Environment, and Health; SD = standard deviation; SFBCS = San Francisco Bay Area Breast Cancer Study
\* Positive family history of breast cancer in first degree relative only
† Positive family history of breast cancer in first or second degree relative
‡ Positive family history of breast cancer in any relative
§ Percentage within cases with known ER status
|| Includes 2 cases with borderline ER status
¶ Calculated for all datasets

1. The San Francisco Bay Area Breast Cancer Study (SFBCS) plus the Northern California Breast Cancer Family Registry (NC-BCFR), a population-based case-control study recruiting from the San Francisco Bay Area [22, 23].
2. The Kaiser Permanente Research Project on Genes, Environment, and Health (RPGEH), a biobank recruiting from Northern California and the Pacific Northwest [24].
3. The Multiethnic Cohort (MEC) study, a prospective cohort study recruiting from Southern California and Hawaii [25].
4. The Cancer de Mama (CAMA) study, a population-based case-control study in Mexico [26].
5. The Post-Columbian Study of Environmental and Heritable Causes of Breast Cancer (COLUMBUS-Colombia), a population-based case-control study in southern Colombia [20].
6. The Post-Columbian Study of Environmental and Heritable Causes of Breast Cancer (COLUMBUS-Mexico), a population-based case-control study in Mexico [20]. The COLUMBUS substudies (Colombia and Mexico) were analyzed as separate datasets given differences in study populations and genotyping methods.
7. The Peru Genetics and Genomics of Breast Cancer Study (PEGEN-BC), a case-series from a Peruvian cancer center. Unrelated Peruvian individuals from 1000 Genomes [27] were used as controls.
8. The City of Hope Clinical Cancer Genetics Community Research Network (COH/CCGCRN), the Southern California site of a multisite cancer center and community-based registry for familial breast cancer [28].

All studies obtained local institutional review board approval and written informed consent from participants.

### Genotyping and genetic ancestry

For all studies except COH/CCGCRN, genotyping was performed using high-density arrays (**Table S1**). Genotyping of COH/CCGCRN was performed using next-generation sequencing with a targeted capture kit that included all 89 SNPs identified as of 2016, prior to publication of the OncoArray GWAS results [3]. Further information about genotyping is provided in the **Supplementary Methods**.

We estimated genetic ancestry from genome-wide markers using the program ADMIXTURE [29] in unsupervised mode with a model containing 4 ancestral populations: European, Indigenous American (IA), African, and East Asian. We used genotype data from 90 European Americans (CEU) and 90 Nigerian Yorubans (YRI) from HapMap [30] to represent European and African populations, respectively. We also included a subset of 504 East Asian individuals from 1000 Genomes [27] and 71 Indigenous Americans previously genotyped on the Affymetrix Axiom LAT1 array [31, 32]. Women with >75% East Asian ancestry were excluded.

### Polygenic risk score

We used a 180-SNP PRS for our primary analysis (**Table S2**). We considered for inclusion 184 SNPs associated with invasive breast cancer with genome-wide significance (p < 5 × 10^−8^) in previous studies [1–4]. These included 172 SNPs from the discovery (n = 65) and replication (n = 107) phases of the Breast Cancer Association Consortium OncoArray study [3], which took place in European (discovery and replication) and Asian (replication only) populations. These SNPs also included nine non-overlapping from GWAS of ER-negative breast cancer [3] and three SNPs from 6q25 discovered in GWAS (rs140068312) [20] and fine-mapping studies (rs3778609, rs851984) [21] in Latinas. Of these 184 SNPs, one pair (rs35054928 and rs2981578) was in linkage disequilibrium (LD) using an r^2^ cutoff of 0.3, and we excluded the latter based on a lower beta coefficient with breast cancer. We also excluded rs17879961 given that it was not polymorphic in our study, and rs2016394 and rs554219 due to a missing call rate > 5%. We included all SNPs regardless of imputation quality, given there were no substantive differences in the associations with breast cancer between the 180-SNP PRS and PRSs constructed with imputation r^2^ thresholds of 0.5 and 0.8, respectively (**Table S3**).

Since targeted genotyping was performed within COH/CCGCRN, genotypes were available for 89 SNPs. We dropped one SNP due to missingness. Of the remaining 88 SNPs, 63 overlapped and 8 had LD proxies (r^2^ > 0.7) with the 180 SNPs comprising the main PRS. We used these 71 SNPs to construct a PRS within COH/CCGCRN. We then constructed a 71-SNP PRS in the 7 remaining datasets using the 63 shared SNPs and 8 respective LD proxies, and pooled all 8 datasets to evaluate the performance of the 71-SNP PRS.

We constructed the PRS as previously described [7, 33]. Briefly, the PRS represents the product of the likelihood ratios across multiple SNPs, assuming each SNP has an independent effect. The likelihood ratio for each SNP was calculated based on the number of risk alleles present and the allele frequency and odds ratio (OR) of the risk allele. We used risk-allele frequencies derived from the Latin American (AMR) population in 1000 Genomes [27] and published ORs for overall breast cancer [3]. The latter predominantly reflects the effect of the SNP within a European population, except for those discovered in Latina studies (**Table S2**) [20, 21].

### Statistical analysis

First, we tested the associations between individual SNPs and breast cancer risk using multivariable logistic regression models adjusted for genetic ancestry and study. Using METAL [34], we performed inverse variance-based meta-analysis of 180 SNPs across 3 studies: COLUMBUS-Colombia, COLUMBUS-Mexico, and pooled SFBCS/NC-BCFR, Kaiser RPGEH, MEC, CAMA, and PEGEN-BC studies.

Next, we tested the crude and adjusted associations between the PRS with breast cancer. Given that genetic ancestry and study were possible confounders of this association (Tables 2 **and S4**), we adjusted for both in our main analysis. To do so, we performed linear regression of study and ancestry on the PRS (dependent variable). We then used the residual as the main predictor in univariate logistic regression with breast cancer as the outcome. We analyzed the residual as a continuous variable normalized to the mean and standard deviation (SD) in controls. We tested the discrimination of the adjusted PRS by estimating the area under the receiver operating characteristic curve (AUROC). We tested calibration using the Hosmer-Lemeshow test across deciles of the adjusted PRS, with the 40-50^th^ and 50-60^th^ deciles combined and used as the reference group.

**Table 2.** Association between 180-SNP and 71-SNP PRS and breast cancer risk

|  | 180-SNP PRS* |  |  |  | 71-SNP PRS† |  |  |  |
| --- | --- | --- | --- | --- | --- | --- | --- | --- |
|  | Controls | Cases | OR (95% CI)‡ | P-trend§ | Controls | Cases | OR (95% CI)‡ | P-trend§ |
| Continuous PRS (per SD) | 7622 | 4658 | 1.58 (1.52 to 1.64) |  | 7927 | 5697 | 1.51 (1.46 to 1.56) |  |
| Percentiles of PRS |  |  |  | <0.001 |  |  |  | <0.001 |
| <10 | 762 | 196 | 0.46 (0.39 to 0.55) |  | 793 | 278 | 0.54 (0.47 to 0.64) |  |
| 10-20 | 763 | 223 | 0.52 (0.44 to 0.62) |  | 792 | 345 | 0.68 (0.58 to 0.79) |  |
| 20-30 | 762 | 340 | 0.80 (0.69 to 0.93) |  | 793 | 379 | 0.74 (0.64 to 0.86) |  |
| 30-40 | 761 | 335 | 0.79 (0.68 to 0.92) |  | 792 | 430 | 0.84 (0.73 to 0.97) |  |
| 40-60 | 1525 | 850 | 1 (referent) |  | 1587 | 1021 | 1 (referent) |  |
| 60-70 | 763 | 498 | 1.17 (1.02 to 1.35) |  | 791 | 656 | 1.29 (1.13 to 1.47) |  |
| 70-80 | 762 | 593 | 1.40 (1.22 to 1.60) |  | 793 | 694 | 1.36 (1.20 to 1.55) |  |
| 80-90 | 761 | 728 | 1.72 (1.50 to 1.96) |  | 793 | 832 | 1.63 (1.44 to 1.85) |  |
| >90 | 763 | 895 | 2.10 (1.85 to 2.39) |  | 793 | 1062 | 2.08 (1.84 to 2.35) |  |

Abbreviations: CI = confidence interval; OR = odds ratio; PRS = polygenic risk score; SD = standard deviation
\* Calculated in case-control analysis in 7 datasets, excluding COH/CCGCRN data (n = 12,280)
† Calculated in case-control analysis of all datasets (n = 13,624)
‡ Odds ratio from multivariable logistic regression of PRS adjusted for study and genetic ancestry
§ P-value for test of linear trend between per-decile estimates

To examine the ancestry-specific performance of the PRS, we divided the pooled dataset into quartiles of IA ancestry. We performed logistic regression within each quartile of IA ancestry and compared the resulting coefficients using a Wald test of linear hypothesis. To compare AUROC estimates, we performed a test of equality of AUROC as described by DeLong [35]. Given differences in the population structures between U.S. Latina and Latin-American studies, we also examined ancestry-specific performance of the PRS by geographic origin of study, specifically U.S. (SFBCS/NC-BCFR, RPGEH, MEC) versus Latin-American (CAMA, COLUMBUS, PEGEN-BC).

All tests for significance used two-sided alpha = 0.05. We developed the script to calculate the PRS using R (The R Foundation). We performed all statistical analyses using Stata 14.2 (StataCorp, College Station, TX).

## Results

### Study characteristics

Our pooled data included 13,624 women from 8 studies, for a total of 5,697 cases and 7,927 controls (Table 1). Across all studies, ancestry was predominantly European and Indigenous American (IA). There was substantial variation in ancestry within and across studies (**Supplementary Figure S2**). For instance, PEGEN-BC in Peru had the highest average IA ancestry (76% in cases and controls) while RPGEH in Northern California had the lowest (27% in cases, 29% in controls). Within each study, cases tended to have similar or lower IA ancestry than controls, as previously reported [36, 37]. In the pooled analysis, cases had higher IA ancestry since nearly half the controls came from RPGEH, the study with the lowest IA ancestry.

### Association of PRS with breast cancer risk

We first examined the associations between individual SNPs and breast cancer risk. Of 180 SNPs, 142 had associations that were directionally consistent with those reported in European populations (**Table S2**) [3]. Forty-four SNPs were nominally significant (p < 0.05), with 39 also directionally consistent. Six SNPs remained significant to p < 2.8×10^−4^ after Bonferroni correction for multiple testing. Nineteen SNPs displayed heterogeneous associations across studies (P_het_ < 0.05). For both PRSs, the mean unadjusted PRS was higher in cases than controls (Table 1, **Figure S1**).

Our main analysis evaluated the performance of a 180-SNP PRS in 12,280 women (4,658 cases and 7,622 controls) from 7 studies, excluding COH/CCGCRN given that only 89 SNPs were genotyped in that study. The unadjusted 180-SNP PRS was strongly associated with breast cancer risk, OR per SD increment = 1.70 (95% CI 1.63 to 1.78). Adjusting for genetic ancestry and study slightly attenuated the association, OR = 1.58 (95% CI 1.52 to 1.64) (Table 2). The associations with breast cancer risk were especially pronounced among extremes of the PRS. Compared with women with a PRS in the 40-60^th^ percentile, women with a PRS in the bottom decile had an OR of 0.46 (95% CI 0.39 to 0.55), whereas those with a PRS in the top decile had an OR of 2.10 (95% 1.85 to 2.39). The AUROC for the 180-SNP PRS was 0.63 (95% CI 0.62 to 0.64), Figure 1A. The Hosmer-Lemeshow test suggested good fit, χ^2^ = 10.45 (p = 0.32), Figure 2A.

**Figure 1.**
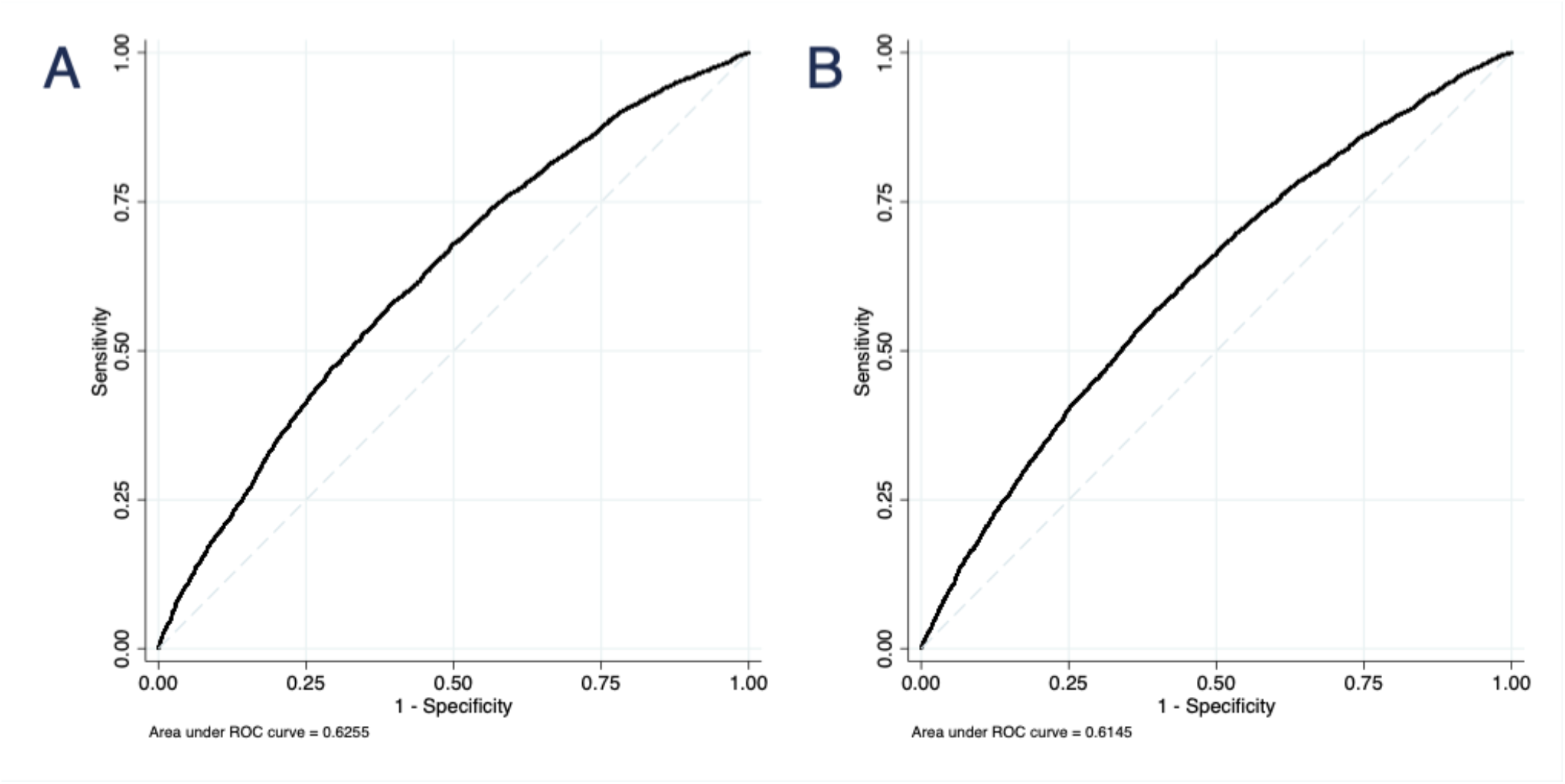
Receiver operating characteristic curves for two polygenic risk scores. The 180-SNP PRS (A) had AUROC = 0.63 (95% CI 0.62 to 0.64) in 7 datasets, excluding COH/CCGCRN (n = 12,280). The 71-SNP PRS (B) had AUROC = 0.61 (95% CI 0.61 to 0.62) in all datasets (n = 13,624).

**Figure 2.**
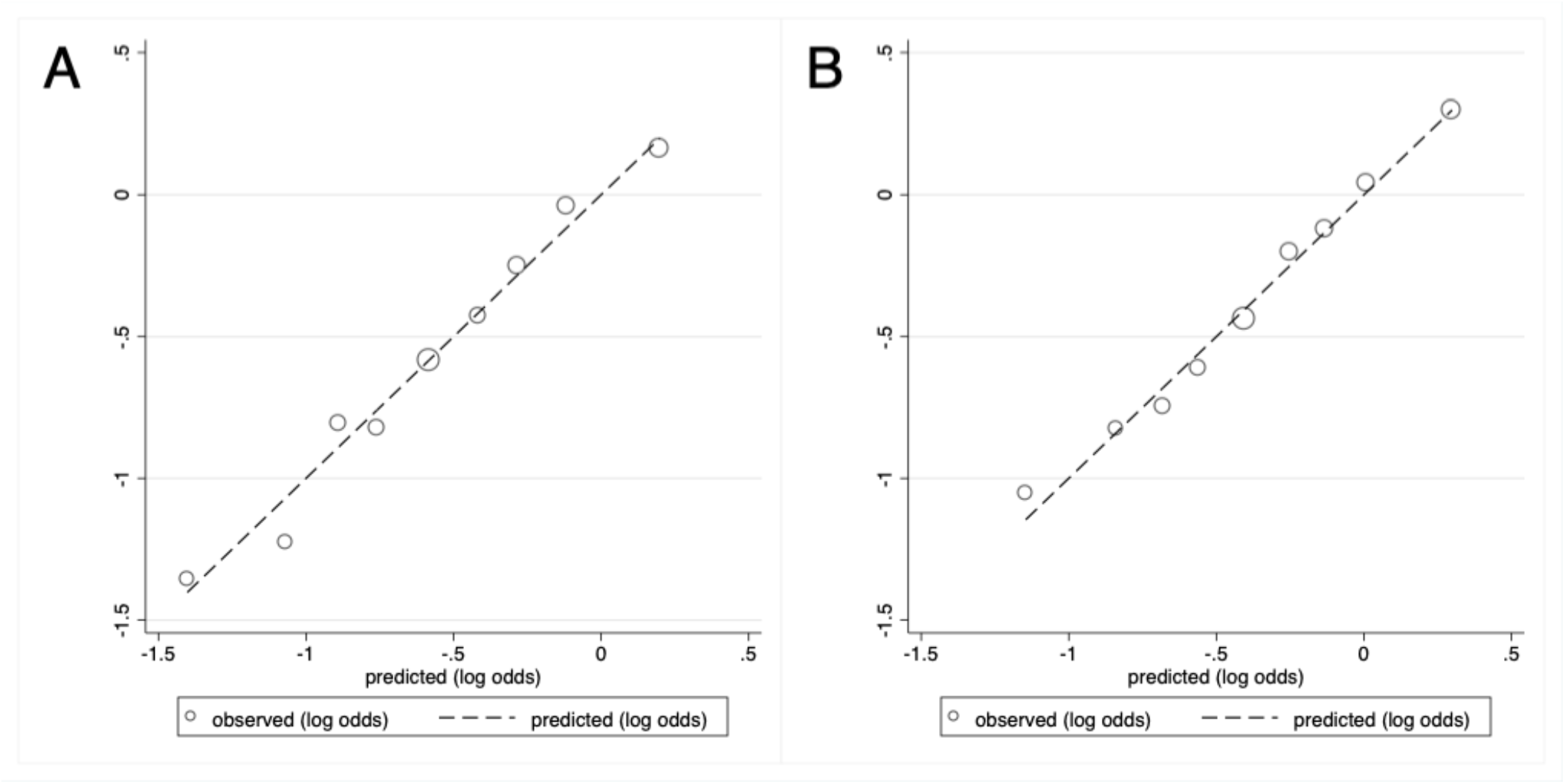
Calibration plots for: (A) the 180-SNP PRS in 7 datasets, excluding COH/CCGCRN (n = 12,280) and (B) the 71-SNP PRS (B) in all datasets (n = 13,624). Graph depicts predicted versus observed proportions of cases within each decile of the log-normalized PRS. Each circle corresponds to a decile of the PRS, with the middle (largest) circle representing the 40-60^th^ percentile. Hosmer-Lemeshow p-value = 0.32 for 180-SNP PRS and 0.68 for 71-SNP PRS.

Our secondary analysis evaluated the performance of a 71-SNP PRS in 13,624 women (5,697 cases and 7,927 controls) from 8 studies, including COH/CCGCRN. Compared with the 180-SNP PRS, the unadjusted 71-SNP PRS had a similar association with breast cancer risk (OR = 1.70, 95% CI 1.62 to 1.79), although adjusting for study and genetic ancestry resulted in larger attenuation of its effect (Table 2, Figure 1B). The discrimination of the 71-SNP PRS was slightly lower and the Hosmer-Lemeshow test was again suggestive of good fit, χ^2^ = 6.59 (p = 0.68), Figure 2B. To assess whether inclusion of COH/CCGCRN participants affected these associations, we tested the 71-SNP PRS with COH/CCGCRN excluded and found similar results (**Table S5**).

### Performance of PRS by Indigenous American ancestry

The 180-SNP PRS displayed similar performance regardless of IA ancestry, with comparable ORs and AUROCs across quartiles of IA ancestry (Table 3). In contrast, the 71-SNP PRS performed worse in the top compared to the bottom quartile, [OR 1.46 (95% CI 1.36 to 1.56) vs OR 1.68 (95% CI 1.55 to 1.83), p = 0.01]. This corresponded to top versus bottom quartile AUROCs of 0.61 (95% CI 0.59 to 0.63) and 0.64 (95% CI 0.62 to 0.66), respectively (p = 0.02). Given differences in ancestry structure between U.S. Latinas and Latin-American women, we stratified the analysis by geographic origin of study. Among 7,317 women from the U.S. studies, the 180-SNP PRS performed best in the bottom quartile of IA ancestry (**Table S6**). However, among the 4,963 women from the Latin-American studies, the 180-SNP PRS performed similarly across quartiles of IA ancestry (**Table S6**).

**Table 3.**
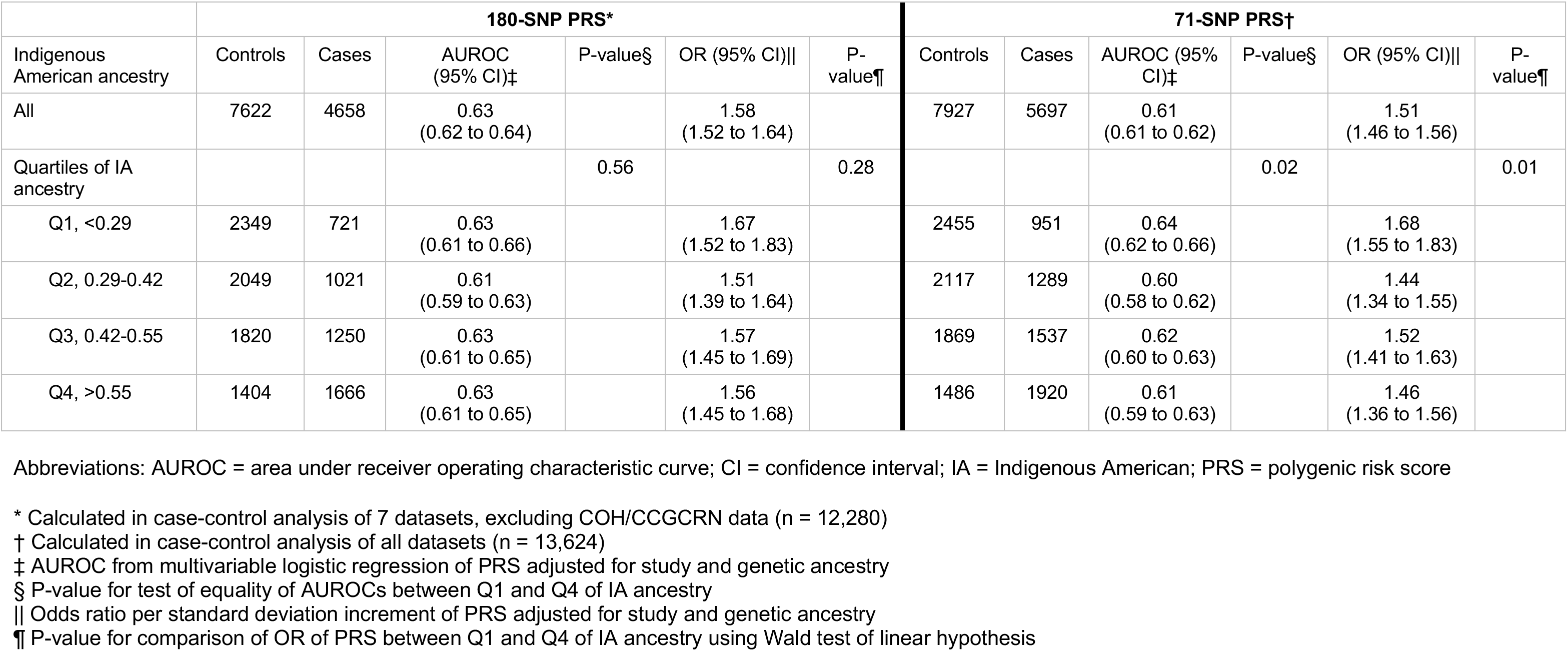
Area under the receiver operating characteristic curve and odds ratios per standard deviation of the 71-SNP PRS and 180-SNP PRS in Hispanics, by quartiles of Indigenous American ancestry

|  | 180-SNP PRS* |  |  |  |  |  | 71-SNP PRS† |  |  |  |  |  |
| --- | --- | --- | --- | --- | --- | --- | --- | --- | --- | --- | --- | --- |
| Indigenous American ancestry | Controls | Cases | AUROC (95% CI)‡ | P-value§ | OR (95% CI)\|\| | P-value¶ | Controls | Cases | AUROC (95% CI)‡ | P-value§ | OR (95% CI)\|\| | P-value¶ |
| All | 7622 | 4658 | 0.63<br>(0.62 to 0.64) |  | 1.58<br>(1.52 to 1.64) |  | 7927 | 5697 | 0.61<br>(0.61 to 0.62) |  | 1.51<br>(1.46 to 1.56) |  |
| Quartiles of IA ancestry |  |  |  | 0.56 |  | 0.28 |  |  |  | 0.02 |  | 0.01 |
| Q1, <0.29 | 2349 | 721 | 0.63<br>(0.61 to 0.66) |  | 1.67<br>(1.52 to 1.83) |  | 2455 | 951 | 0.64<br>(0.62 to 0.66) |  | 1.68<br>(1.55 to 1.83) |  |
| Q2, 0.29-0.42 | 2049 | 1021 | 0.61<br>(0.59 to 0.63) |  | 1.51<br>(1.39 to 1.64) |  | 2117 | 1289 | 0.60<br>(0.58 to 0.62) |  | 1.44<br>(1.34 to 1.55) |  |
| Q3, 0.42-0.55 | 1820 | 1250 | 0.63<br>(0.61 to 0.65) |  | 1.57<br>(1.45 to 1.69) |  | 1869 | 1537 | 0.62<br>(0.60 to 0.63) |  | 1.52<br>(1.41 to 1.63) |  |
| Q4, >0.55 | 1404 | 1666 | 0.63<br>(0.61 to 0.65) |  | 1.56<br>(1.45 to 1.68) |  | 1486 | 1920 | 0.61<br>(0.59 to 0.63) |  | 1.46<br>(1.36 to 1.56) |  |

Abbreviations: AUROC = area under receiver operating characteristic curve; CI = confidence interval; IA = Indigenous American; PRS = polygenic risk score
\* Calculated in case-control analysis of 7 datasets, excluding COH/CCGCRN data (n = 12,280)
† Calculated in case-control analysis of all datasets (n = 13,624)
‡ AUROC from multivariable logistic regression of PRS adjusted for study and genetic ancestry
§ P-value for test of equality of AUROCs between Q1 and Q4 of IA ancestry
|| Odds ratio per standard deviation increment of PRS adjusted for study and genetic ancestry
¶ P-value for comparison of OR of PRS between Q1 and Q4 of IA ancestry using Wald test of linear hypothesis

## Discussion

We found that PRSs primarily consisting of SNPs identified in European populations were predictive of breast cancer risk in Latinas. Our 180-SNP PRS had an adjusted OR per SD increment of 1.58 (95% CI 1.52 to 1.64) and an AUROC of 0.63 (95% CI 0.62 to 0.64). These results are comparable to those of European studies, which tested PRSs including 77 to 3820 SNPs and reported ORs per SD between 1.46-1.66 and AUROCs between 0.60-0.64 [5, 38]. Our 71-SNP PRS performed worse than the 180-SNP PRS, though the difference was modest.

Ours is the largest study to date on breast cancer PRS in Latinas and extends the literature by refining estimates of PRS performance in this population. Allman, et al [14] reported that a 71-SNP PRS had an OR per SD increment of 1.39 (95% CI 1.18 to 1.64) and AUROC of 0.59 (95% CI 0.54 to 0.64) among U.S. Latinas, but this study included only 147 cases and did not account for ancestry.

We could not definitively determine whether PRS performance varies by ancestry. Differential PRS performance by genetic ancestry might be expected given differences in LD structures between European and non-European populations, which can attenuate the associations between GWAS hits discovered in Europeans and causal SNPs in LD. Additionally, causal alleles may only be present in certain populations. However, the 180-SNP PRS performed similarly across quartiles of IA ancestry. In contrast, the 71-SNP PRS performed better in the bottom quartile of IA ancestry, corresponding to higher European ancestry. This analysis included 1,039 additional cases from COH/CCGCRN and may have had greater statistical power to detect differences in performance by IA ancestry.

A major strength of our study was the size and diversity of our study population. Additionally, we accounted for genetic ancestry, which can bias associations in genetic studies [39]. Given that ancestry was a confounder and an independent predictor of breast cancer risk, we used a novel approach to calculate an “ancestry-adjusted” PRS. We also examined PRS performance by IA ancestry, which has not been previously done. Another strength was the inclusion of several large, diverse breast cancer studies representing populations from several geographic areas (Western U.S., Central and South America) and including women with varying degrees of IA versus European ancestry.

Our results should be interpreted in light of three limitations. First, the generalizability of our findings is limited to Latina populations with similar distributions of genetic ancestry. Although the ancestry composition of our study resembled that of other large studies of Latinas from the western U.S. and Central/South America [19, 40], our results may not be generalizable to Caribbean Latinas, whose population structures have higher proportions of African ancestry [17–19]. We did not test the performance of PRS according to African ancestry given that our study population was predominantly Latinas with limited African ancestry. Secondly, our analysis included women from community-based and familial breast cancer clinics and may include moderate or high-penetrance mutation carriers. While PRS is associated with breast cancer risk in mutation carriers and women with elevated familial risk, the magnitudes of these associations vary slightly from those in the average-risk population [41]. Finally, we tested a PRS containing 180 SNPs representing known GWAS hits at the time of analysis. However, others have constructed PRSs comprising 313 and 3820 SNPs by including SNPs that did not have genome-wide significant associations with breast cancer [38]. Though these expanded PRSs performed better than a 77-SNP PRS, there was little difference in performance between the 313-SNP and 3820-SNP PRSs [38]. We included only SNPs with genome-wide significant associations in our PRS since these signals may be more robust across ancestry. The AUROC for our 180-SNP PRS (0.63) was similar to that of the 313-SNP PRS [38].

Our results suggest that the PRS has predictive value in Latinas, a large and rapidly-growing population in the U.S. Although studies on the ability of the PRS to inform decisions around screening and prevention are underway [9], several commercial genetic testing laboratories already return PRS results to women of European descent who tested negative for deleterious mutations [10, 11]. If this practice were extended to Latinas, one could expect the PRS to perform comparably well. Even if the performance of the PRS were slightly attenuated in Latinas of higher Indigenous American ancestry, this should not preclude its use in this population. Instead, results could account for this attenuation and model the joint effects of PRS and ancestry.

Though our findings suggest that the PRS can predict breast cancer risk in Latinas, they do not nullify the prospect of disparities in genetic discovery research [42, 43]. Whereas we studied mostly common variants, rare variants display more geographic clustering [44]. As genetic association studies identify more rare variants, those discovered in European populations will be less generalizable to other populations. Thus, high-quality genetic studies in non-European populations remain a priority. Fine-mapping in large datasets may enhance the identification of causal SNPs associated with breast cancer risk. Likewise, GWAS should be intentional about including Latinas, particularly those with higher IA and/or African ancestry. In addition, future studies should prospectively assess prediction and examine the contribution of PRS to clinical risk models. Though one such trial is currently using the PRS to tailor decision-making around breast cancer screening and prevention [9], similar clinical effectiveness studies also should aim to recruit diverse women.

## Notes

We thank the participants of the SFBCS/NC-BCFR, Kaiser Permanente RPGEH, MEC, CAMA, COLUMBUS, PEGEN-BC, and COH-CCGCRN studies. The authors declare no competing interests.

The contributors from the COLUMBUS Consortium (in alphabetical order) include: Jennyfer Benavides (Universidad del Tolima, Ibagué, Colombia), Mabel Bohorquez (Universidad del Tolima, Ibagué, Colombia), Fernando Bolaños (Hospital Hernando Moncaleano Perdomo, Neiva, Colombia), Luis G Carvajal-Carmona (Universidad del Tolima, Ibagué, Colombia, University of California Comprehensive Cancer Center, Sacramento, USA, Fundación de Genética y Genómica, Medellín, Colombia, Genome Center and Department of Biochemistry and Molecular Medicine, University of California, Davis, Davis, CA, USA), Jenny Carmona (Dinámica IPS, Medellín, Colombia), Ángel Criollo (Universidad del Tolima, Ibagué, Colombia), Magdalena Echeverry (Universidad del Tolima, Ibagué, Colombia), Ana Estrada (Universidad del Tolima, Ibagué, Colombia), Gilbert Mateus (Hospital Federico Lleras Acosta, Ibagué, Colombia), Raúl Murillo (Pontificia Universidad Javeriana, Bogotá, Colombia), Justo Ramirez (Hospital Hernando Moncaleano Perdomo, Neiva, Colombia), Yesid Sánchez (Universidad del Tolima, Ibagué, Colombia), Carolina Sanabria (Instituto Nacional de Cancerología, Bogotá, Colombia), Martha Lucia Serrano (Instituto Nacional de Cancerología, Bogotá, Colombia), John Jairo Suarez (Universidad del Tolima, Ibagué, Colombia), Alejandro Vélez (Dinámica IPS, Medellín, Colombia, Hospital Pablo Tobón Uribe, Medellín, Colombia).

