## Supplementary Materials for "A polygenic risk score for breast cancer in U.S. Latinas and Latin-American women"

**Affiliations**
1. Division of General Internal Medicine, Department of Medicine, University of California, San Francisco, San Francisco, CA, USA
2. UC Davis Genome Center, University of California, Davis, Davis, CA, USA
3. School of Public Health, University of California, Berkeley; Berkeley, CA, USA
4. Natera, San Carlos, CA, USA
5. Unidad de Investigación en Enfermedades Infecciosas, Instituto Mexicano del Seguro Social; Mexico City, Mexico
6. Grupo de Citogenética, Filogenia y Evolución de Poblaciones, Facultades de Ciencias y Facultad de Ciencias de la Salud, Universidad del Tolima, Ibagué, Colombia
7. Hospital de Gineco Obstetricia No.4 “LCA”, Instituto Mexicano del Seguro Social; Mexico City, Mexico
8. Department of Preventive Medicine, Norris Comprehensive Cancer Center, Keck School of Medicine, University of Southern California, Los Angeles, CA, USA
9. Department of Medicine, Stanford University School of Medicine, Stanford, CA, USA
10. Stanford Cancer Institute, Stanford University School of Medicine, Stanford, CA, USA
11. Division of Research, Kaiser Permanente Northern California, Oakland, CA, USA
12. Instituto Nacional de Salud Pública, Cuernavaca, Mexico
13. Instituto Nacional de Enfermedades Neoplásicas (INEN), Surquillo, Peru
14. Division of Clinical Genetics, City of Hope National Medical Center, Duarte, CA, USA
15. Instituto Nacional de Enfermedades Neoplásicas (INEN), Lima, Peru
16. Department of Biochemistry and Molecular Medicine, School of Medicine, University of California, Davis, Sacramento, CA, USA
17. Population Science and Health Disparities Program, University of California Davis Comprehensive Cancer Center, Sacramento, CA, USA
18. Department of Population Sciences, Beckman Research Institute of City of Hope, Duarte, CA, USA

**Corresponding author:**
Yiwey Shieh, MD
Division of General Internal Medicine
Department of Medicine, University of California, San Francisco
1545 Divisadero Street, Box 0320
San Francisco, CA 94115 USA

1 (415) 885-7277

**Supplementary Methods**

*Participants*

The San Francisco Bay Area Breast Cancer Study (SFBCS) is a population-based case-control study of breast cancer. The Greater Bay Area Cancer Registry was used to identify breast cancer cases newly diagnosed from 1995-2002 in self-identified Latinas. Random-digit dialing was used to identify controls matched on race/ethnicity and 5-year age group. The Northern California Breast Cancer Family Registry (NC-BCFR) is one of six international sites participating in the Breast Cancer Family Registry. Cases diagnosed from 1995-2009 were identified through the Greater Bay Area Cancer Registry, with controls also frequency matched on race/ethnicity and 5-year age group. We combined the SFBCS and NC-BCFR datasets given that they recruited from the same geographic region and in overlapping time frames. We used KING [1] to identify relative pairs between SFBCS/NC-BCFR using a kinship coefficient of > 0.2. We removed one individual from each relative pair while preferentially keeping cases. We also removed overlapping individuals with RPGEH, yielding a total of 942 cases and 589 controls.

The Kaiser Permanente Northern California Research Project on Genes, Environment, and Health (RPGEH) is a multiracial/multiethnic biobank comprised of consenting members of the Kaiser Permanente Health Plan. A subset of cohort members was genotyped, forming the Genetic Epidemiology Research on Adult health and Aging (GERA) cohort. We obtained genotype data for incident and prevalent cases of breast cancer in Latinas from the GERA cohort (total n = 222) plus non-breast cancer Latina controls (n = 3,563).

The Multiethnic Cohort (MEC) study is a prospective cohort study of cancer that recruited in California (predominantly Los Angeles County) and Hawaii. We analyzed a nested sample of 532 cases, representing all incident cases of breast cancer in women aged 45 and over at diagnosis who self-identified as Latina. We analyzed 1469 female controls matched on age (within 5 years) and self-identified ethnicity.

The Cancer de Mama (CAMA) study is a population-based case-control study of breast cancer in Mexico City, Monterrey, and Veracruz, Mexico. We analyzed 709 cases aged 35-69 years and diagnosed from 2005-2007 at one of eleven hospitals in the study region. We also included 702 controls frequency-matched by 5-year age group.

The Post-Columbian Study of Environmental and Heritable Causes of Breast Cancer (COLUMBUS) is a case-control study of breast cancer with individuals recruited throughout large cancer hospitals in central and southern Colombia and Mexico City. From Colombia, we analyzed 954 cases aged 21-88 years and diagnosed 2011-2017. We included 768 cancer-free controls recruited from the same institutions as cases and matched on education, socioeconomic status, and local origin. Controls did not have a family history of breast cancer in a first-degree relative.

From Mexico City, Mexico, we analyzed data from 481 cases aged 25-89 and diagnosed 2010-2018 at the Social Security Hospital in Mexico City, the largest cancer hospital in the country. As controls, we included 453 blood donors without breast cancer recruited from the same hospital.

The Peru Genetics and Genomics of Breast Cancer Study (PEGEN-BC) is a case-series study. We recruited 1650 participants from the Instituto Nacional de Enfermedades Neoplásicas (INEN) in Lima, Peru. The initial inclusion criteria were an invasive breast cancer diagnosed between the ages of 21-79 years, and after 2010. In actuality, our sample included 79 women diagnosed before 2010 (5 of whom were diagnosed before the year 2000), and 4 women older than 79. A recruiter working locally at INEN identified eligible breast cancer patients who had upcoming appointments with their oncology provider. During appointment check-in, the recruiter approached eligible patients, provided the patients with information about the study, and obtained written informed consent if they showed interest in participating in the study. At the end of their clinic visit, blood was drawn by a certified phlebotomist at the INEN central laboratory. The participants were given a short questionnaire with questions on lifestyle, family history, and reproductive behavior. Clinical data including tumor characteristics, treatment, and disease progression was abstracted from medical records at INEN and from outside clinics, if necessary. The current analysis included 818 breast cancer cases for whom genome-wide genotypes were available. We analyzed 85 unrelated Peruvian samples from 1000 Genomes as controls.

From the City of Hope (COH), we included self-identified Latina participants in the Clinical Cancer Genetics Community Research Network (CCGCRN), a multi-site registry of cancer center and community-based clinics serving individuals with a personal or family history of breast cancer. Cases had a diagnosis of invasive breast cancer. Controls were recruited locally and primarily at community health fairs, with an additional 76 controls from the California Teachers Study [2]. We excluded first-degree relatives of MEC participants. We analyzed 1,039 cases and 305 controls.

*Genotyping*

We performed genotyping using the following arrays (**Table S1**): Affymetrix 6.0 for SFBCS/NC-BCFR samples; Affymetrix LAT for Kaiser Permanente RPGEH samples; Illumina 660 W-Quad for MEC cases and Illumina 2.5M for MEC controls; Illumina OncoArray for CAMA samples; Axiom GW LAT Axiom Custom for Colombian and Mexican controls, Axiom UK Biobank for COLUMBUS Colombian samples, and Affymetrix Axiom Precision Medicine Research Array (PMRA) for COLUMBUS Mexican cases and controls, and for the PEGEN-BC samples.

Sequence results were aligned using Burrows-Wheeler Aligner [3] and genotype calls were made using Genome Analysis Toolkit [4] as previously described [5].

We excluded samples with >5% missing genotypes and SNPs with >5% missingness. All genotype datasets were mapped to Genome Reference Consortium Human Build 37 (hg19). Each dataset (with the exception of COH/CCGCRN) was phased using EAGLE v2.3 [6] and imputed to the Haplotype Reference Consortium reference panel [7] using the Michigan Imputation Server [8]. Since the MEC datasets included both 660K and 2.5M arrays, we performed imputation using the overlapping SNPs (n = 192,795) given that pooling the separately-imputed datasets resulted in false positives.

**Table S1.** Characteristics of 8 studies included in the analysis

| Study | Recruitment area | Controls | Cases | Years of diagnosis (cases) | Age of cases, years | Control group composition | Genotyping Platform | Genotyping site | Imputation Reference |
|---|---|---|---|---|---|---|---|---|---|
| SFBCS/NC-BCFR | San Francisco Bay Area, USA | 589 | 942 | 1995-2002 (SFBCS), 1995-2007 (NC-BCFR) | 35-79 (SFBCS), 18-64 (NC-BCFR) | Identified through random digit-dialing and frequency matched on 5-year age groups (both) | Affymetrix 6.0 | University of California, San Francisco (lab of Esteban Burchard) | 1000 Genomes, HRC |
| Kaiser RPGEH | Northern California, USA | 3563 | 222 | | >18 | Overall cohort of self-identified Latinas without breast cancer | Affymetrix LAT | University of California, San Francisco (Institute of Human Genetics) | 1000 Genomes, HRC |
| MEC | Los Angeles County and Hawaii, USA | 1469 | 532 | | >45 | Frequency matched on 5-year age groups and ethnicity. | Illumina 660 W-Quad (cases), Omni 2.5M (controls) | University of Southern California (cases), Broad Institute (controls) | 1000 Genomes, HRC |
| CAMA | Mexico City, Monterrey, and Veracruz, Mexico | 702 | 709 | 2005-2007 | 35-69 | Frequency matched on 5-year age groups | Illumina OncoArray | Quebec Genome Center | HRC |
| COLUMBUS (Colombia) | Central and Southern Colombia | 761 | 954 | 2011-2017 | 21-88 | Recruited from same geographic region | Affymetrix Axiom GW LAT and Custom (controls), Axiom UKBiobank (cases) | University of North Carolina (cases), bioBANC (controls) | 1000 Genomes, HRC |
| COLUMBUS (Mexico) | Mexico City, Mexico | 453 | 481 | 2010-2018 | 25-89 | Recruited from same geographical region | Affymetrix Axiom PMRA (cases, controls), Axiom GW LAT and Custom (controls) | Affymetrix (cases, controls), bioBANC (controls) | 1000 Genomes, HRC |
| Peru | Lima, Peru | 85 | 818 | 1985-2018 | 22-91 | Unrelated individuals from 1000 Genomes recruited in Lima, Peru | Affymetrix Axiom PMRA (cases) | University of California, San Francisco | 1000 Genomes |
| COH/CCGCRN | Southern California, USA | 305 | 1039 | 1998-present | 18-85 | Recruited from same geographic region | Next-generation sequencing with a targeted capture kit | City of Hope | N/A |

Abbreviations: CAMA = Cancer de Mama; COH/CCGCRN = Clinical Cancer Genetics Community Research Network; COLOMBUS = Colombian Study of Environmental and Heritable Causes of Breast Cancer; HRC = Haplotype Reference Consortium; MEC = Multiethnic Cohort; NCBCFR = Northern California Breast Cancer Family Registry; PRS = polygenic risk score; RPGEH = Research Project on Genes, Environment, and Health; SD = standard deviation; SFBCS = San Francisco Bay Area Breast Cancer Study

**Table S2.** List of SNPs in the 180-SNP PRS with allele frequencies and associations from 7 pooled Hispanic datasets

| | | | | | COLUMBUS - Mexico | | | COLUMBUS - Colombia | | | All other studies† | | | Meta-analysis | | |
|---|---|---|---|---|---|---|---|---|---|---|---|---|---|---|---|---|
| rsid* | chr | pos | Risk | Other | RAF | OR (95% CI) | *P* | RAF | OR (95% CI) | *P* | RAF | OR (95% CI) | *P* | OR (95% CI) | *P* | $P_{het}$ |
| rs616488 | 1 | 10566215 | A | G | 0.48 | 1.11 (0.92-1.35) | 0.28 | 0.57 | 1.12 (0.98-1.29) | 0.10 | 0.56 | 1.08 (1.00-1.17) | 0.05 | 1.09 (1.03-1.16) | 5.8E-03 | 0.88 |
| rs2992756 | 1 | 18807339 | T | C | 0.38 | 0.91 (0.75-1.12) | 0.37 | 0.42 | 1.00 (0.87-1.14) | 0.94 | 0.40 | 0.98 (0.90-1.05) | 0.54 | 0.97 (0.91-1.04) | 0.41 | 0.78 |
| rs4233486 | 1 | 41380440 | T | C | 0.76 | 0.89 (0.70-1.12) | 0.32 | 0.76 | 1.03 (0.88-1.21) | 0.67 | 0.71 | 1.06 (0.98-1.16) | 0.16 | 1.04 (0.97-1.12) | 0.29 | 0.36 |
| rs79724016 | 1 | 42137311 | T | G | 0.99 | 1.78 (0.65-4.85) | 0.26 | 0.98 | 1.24 (0.70-2.19) | 0.46 | 0.98 | 0.99 (0.75-1.31) | 0.94 | 1.07 (0.84-1.36) | 0.60 | 0.46 |
| rs1707302 | 1 | 46600917 | G | A | 0.62 | 0.93 (0.76-1.13) | 0.47 | 0.61 | 1.13 (0.98-1.29) | 0.09 | 0.65 | 1.05 (0.97-1.14) | 0.23 | 1.05 (0.99-1.12) | 0.12 | 0.29 |
| rs140850326 | 1 | 50846032 | CAAAGGGCAAGATCTCCTTTTT | C | 0.27 | 0.99 (0.80-1.24) | 0.94 | 0.42 | 1.05 (0.92-1.21) | 0.45 | 0.37 | 0.98 (0.90-1.06) | 0.62 | 1.00 (0.93-1.07) | 0.96 | 0.66 |
| rs17426269 | 1 | 88156923 | A | G | 0.07 | 0.62 (0.41-0.93) | 0.02 | 0.09 | 1.11 (0.88-1.40) | 0.39 | 0.09 | 1.06 (0.93-1.22) | 0.37 | 1.03 (0.92-1.15) | 0.59 | 0.04 |
| rs11552449 | 1 | 114448389 | T | C | 0.47 | 0.84 (0.69-1.03) | 0.09 | 0.38 | 0.99 (0.86-1.14) | 0.88 | 0.35 | 0.92 (0.85-0.99) | 0.03 | 0.92 (0.86-0.99) | 0.02 | 0.41 |
| rs7529522 | 1 | 118230221 | C | T | 0.44 | 0.98 (0.80-1.20) | 0.85 | 0.40 | 0.98 (0.85-1.12) | 0.77 | 0.37 | 1.06 (0.98-1.14) | 0.16 | 1.03 (0.97-1.10) | 0.34 | 0.56 |
| rs11249433 | 1 | 121280613 | G | A | 0.16 | 0.99 (0.75-1.30) | 0.93 | 0.24 | 1.24 (1.06-1.45) | 6.8E-03 | 0.27 | 1.05 (0.96-1.15) | 0.32 | 1.08 (1.01-1.17) | 0.03 | 0.14 |
| rs12405132 | 1 | 145644984 | C | T | 0.71 | 1.11 (0.91-1.37) | 0.31 | 0.66 | 0.98 (0.85-1.14) | 0.83 | 0.68 | 1.03 (0.95-1.12) | 0.48 | 1.03 (0.96-1.10) | 0.42 | 0.63 |
| rs12048493 | 1 | 149927034 | C | A | 0.22 | 1.00 (0.79-1.27) | 0.97 | 0.27 | 1.13 (0.97-1.31) | 0.12 | 0.23 | 1.02 (0.92-1.12) | 0.75 | 1.04 (0.96-1.13) | 0.29 | 0.50 |
| rs4971059 | 1 | 155148781 | A | G | 0.61 | 1.02 (0.83-1.25) | 0.87 | 0.55 | 0.99 (0.87-1.14) | 0.91 | 0.50 | 1.05 (0.97-1.14) | 0.21 | 1.03 (0.97-1.10) | 0.30 | 0.76 |
| rs35383942 | 1 | 201437832 | T | C | 0.03 | 0.76 (0.38-1.53) | 0.44 | 0.02 | 1.25 (0.83-1.89) | 0.29 | 0.03 | 0.99 (0.78-1.25) | 0.91 | 1.02 (0.84-1.24) | 0.84 | 0.43 |
| rs6678914 | 1 | 202187176 | G | A | 0.77 | 0.90 (0.72-1.14) | 0.40 | 0.70 | 1.08 (0.93-1.26) | 0.33 | 0.70 | 1.10 (1.01-1.20) | 0.03 | 1.08 (1.00-1.15) | 0.04 | 0.30 |
| rs4951011 | 1 | 203766331 | G | A | 0.39 | 1.14 (0.94-1.39) | 0.19 | 0.35 | 1.05 (0.91-1.21) | 0.52 | 0.30 | 1.05 (0.97-1.14) | 0.23 | 1.06 (0.99-1.13) | 0.09 | 0.74 |
| rs4245739 | 1 | 204518842 | C | A | 0.24 | 1.07 (0.86-1.34) | 0.54 | 0.27 | 1.01 (0.87-1.19) | 0.86 | 0.26 | 1.03 (0.94-1.12) | 0.56 | 1.03 (0.96-1.10) | 0.45 | 0.92 |
| rs11117758 | 1 | 217220574 | G | A | 0.75 | 0.83 (0.66-1.04) | 0.11 | 0.75 | 1.04 (0.89-1.22) | 0.62 | 0.76 | 1.04 (0.96-1.14) | 0.34 | 1.02 (0.95-1.09) | 0.61 | 0.18 |
| rs72755295 | 1 | 242034263 | G | A | 0.01 | 0.36 (0.07-1.92) | 0.23 | 0.01 | 0.99 (0.39-2.52) | 0.98 | 0.01 | 1.08 (0.69-1.70) | 0.74 | 1.00 (0.67-1.49) | 1.00 | 0.46 |
| rs113577745 | 2 | 10135681 | G | C | 0.04 | 1.11 (0.70-1.77) | 0.64 | 0.05 | 1.03 (0.76-1.40) | 0.84 | 0.07 | 1.10 (0.94-1.28) | 0.25 | 1.09 (0.95-1.24) | 0.23 | 0.93 |
| rs12710696 | 2 | 19320803 | T | C | 0.27 | 1.11 (0.90-1.38) | 0.33 | 0.33 | 0.95 (0.82-1.10) | 0.47 | 0.32 | 1.06 (0.98-1.15) | 0.17 | 1.04 (0.97-1.11) | 0.27 | 0.35 |
| rs200648189 | 2 | 24739694 | CT | C | 0.86 | 1.16 (0.88-1.53) | 0.29 | 0.82 | 1.13 (0.94-1.34) | 0.19 | 0.85 | 1.04 (0.93-1.16) | 0.49 | 1.07 (0.98-1.17) | 0.12 | 0.63 |
| rs6725517 | 2 | 25129473 | A | G | 0.75 | 1.02 (0.81-1.27) | 0.88 | 0.71 | 1.13 (0.97-1.32) | 0.12 | 0.68 | 1.08 (0.99-1.17) | 0.09 | 1.08 (1.01-1.16) | 0.03 | 0.74 |
| rs4577244 | 2 | 29120733 | T | C | 0.62 | 1.05 (0.86-1.29) | 0.61 | 0.45 | 0.89 (0.78-1.02) | 0.11 | 0.48 | 1.03 (0.95-1.11) | 0.45 | 1.00 (0.94-1.07) | 0.97 | 0.18 |
| rs71801447 | 2 | 111925731 | C | CTTATGTT | 0.05 | 1.15 (0.72-1.81) | 0.56 | 0.06 | 0.99 (0.74-1.31) | 0.92 | 0.07 | 1.06 (0.91-1.23) | 0.46 | 1.05 (0.92-1.19) | 0.45 | 0.85 |
| rs4849887 | 2 | 121245122 | C | T | 0.90 | 0.88 (0.63-1.22) | 0.43 | 0.88 | 1.19 (0.96-1.46) | 0.11 | 0.88 | 1.01 (0.90-1.14) | 0.82 | 1.04 (0.94-1.14) | 0.48 | 0.25 |
| rs2016394 | 2 | 172972971 | G | A | 0.55 | 0.95 (0.79-1.16) | 0.64 | 0.52 | 1.03 (0.90-1.18) | 0.66 | 0.56 | 1.13 (1.05-1.22) | 1.4E-03 | 1.09 (1.02-1.16) | 7.1E-03 | 0.19 |
| rs1550623 | 2 | 174212894 | A | G | 0.74 | 0.93 (0.75-1.17) | 0.55 | 0.77 | 1.23 (1.04-1.45) | 0.02 | 0.78 | 1.02 (0.93-1.12) | 0.70 | 1.05 (0.97-1.13) | 0.22 | 0.08 |
| rs1830298 | 2 | 202181247 | C | T | 0.47 | 1.12 (0.93-1.36) | 0.23 | 0.48 | 0.97 (0.84-1.11) | 0.62 | 0.42 | 1.02 (0.95-1.10) | 0.58 | 1.02 (0.96-1.09) | 0.53 | 0.45 |
| rs4442975 | 2 | 217920769 | G | T | 0.27 | 1.03 (0.83-1.29) | 0.76 | 0.36 | 1.08 (0.94-1.25) | 0.26 | 0.36 | 1.16 (1.07-1.26) | 2.5E-04 | 1.13 (1.06-1.21) | 2.6E-04 | 0.49 |
| rs34005590 | 2 | 217963060 | C | A | 0.99 | 0.61 (0.23-1.65) | 0.33 | 0.99 | 1.44 (0.78-2.66) | 0.25 | 0.98 | 1.14 (0.87-1.48) | 0.35 | 1.14 (0.90-1.44) | 0.30 | 0.35 |
| rs16857609 | 2 | 218296508 | T | C | 0.44 | 1.14 (0.94-1.39) | 0.20 | 0.34 | 1.03 (0.89-1.18) | 0.71 | 0.38 | 1.03 (0.95-1.11) | 0.50 | 1.04 (0.97-1.11) | 0.25 | 0.62 |
| rs12479355 | 2 | 227226952 | A | G | 0.72 | 0.92 (0.74-1.14) | 0.44 | 0.74 | 1.03 (0.88-1.20) | 0.75 | 0.76 | 1.02 (0.94-1.12) | 0.58 | 1.01 (0.94-1.09) | 0.72 | 0.65 |
| rs6762644 | 3 | 4742276 | G | A | 0.22 | 0.99 (0.78-1.25) | 0.93 | 0.29 | 1.10 (0.95-1.27) | 0.20 | 0.26 | 1.04 (0.95-1.13) | 0.38 | 1.05 (0.98-1.13) | 0.19 | 0.71 |
| rs4973768 | 3 | 27416013 | T | C | 0.57 | 0.88 (0.72-1.06) | 0.18 | 0.59 | 1.20 (1.04-1.38) | 0.01 | 0.56 | 1.06 (0.98-1.14) | 0.13 | 1.06 (1.00-1.13) | 0.05 | 0.04 |
| rs12493607 | 3 | 30682939 | C | G | 0.35 | 0.98 (0.80-1.20) | 0.87 | 0.24 | 1.12 (0.95-1.31) | 0.17 | 0.33 | 1.00 (0.92-1.08) | 1.00 | 1.02 (0.95-1.09) | 0.61 | 0.44 |
| rs6796502 | 3 | 46866866 | G | A | 0.71 | 1.04 (0.84-1.29) | 0.73 | 0.79 | 1.02 (0.86-1.21) | 0.83 | 0.79 | 0.99 (0.90-1.08) | 0.75 | 1.00 (0.93-1.07) | 0.96 | 0.88 |
| rs1053338 | 3 | 63967900 | G | A | 0.23 | 0.88 (0.70-1.11) | 0.29 | 0.21 | 0.90 (0.76-1.07) | 0.23 | 0.19 | 1.00 (0.91-1.10) | 0.98 | 0.96 (0.89-1.04) | 0.36 | 0.43 |
| rs6805189 | 3 | 71532113 | T | C | 0.83 | 1.05 (0.80-1.38) | 0.73 | 0.71 | 1.10 (0.95-1.27) | 0.22 | 0.71 | 0.93 (0.86-1.02) | 0.14 | 0.98 (0.91-1.06) | 0.60 | 0.17 |

| | | | | | | | | | | | | | | | | |
|---|---|---|---|---|---|---|---|---|---|---|---|---|---|---|---|---|
| rs13066793 | 3 | 87037543 | A | G | 0.97 | 1.13 (0.66-1.93) | 0.66 | 0.93 | 1.18 (0.90-1.55) | 0.23 | 0.95 | 1.00 (0.83-1.20) | 0.98 | 1.06 (0.91-1.23) | 0.45 | 0.58 |
| rs9833888 | 3 | 99723580 | T | G | 0.22 | 1.12 (0.89-1.41) | 0.33 | 0.23 | 1.02 (0.87-1.19) | 0.84 | 0.23 | 1.07 (0.98-1.17) | 0.11 | 1.07 (0.99-1.14) | 0.09 | 0.77 |
| rs34207738 | 3 | 141112859 | C | CTT | 0.19 | 1.53 (1.20-1.95) | 5.8E-04 | 0.27 | 1.12 (0.96-1.30) | 0.14 | 0.27 | 1.03 (0.95-1.13) | 0.46 | 1.09 (1.01-1.17) | 0.02 | 0.01 |
| rs58058861 | 3 | 172285237 | A | G | 0.28 | 0.90 (0.73-1.12) | 0.36 | 0.09 | 1.40 (1.13-1.74) | 2.3E-03 | 0.23 | 1.00 (0.92-1.10) | 0.93 | 1.03 (0.96-1.11) | 0.41 | 0.01 |
| rs6815814 | 4 | 38816338 | C | A | 0.33 | 1.08 (0.87-1.32) | 0.49 | 0.37 | 1.05 (0.91-1.21) | 0.50 | 0.37 | 0.95 (0.88-1.02) | 0.17 | 0.98 (0.92-1.05) | 0.54 | 0.29 |
| chr4:84370124 | 4 | 84370124 | TAA | TA | 0.30 | 1.06 (0.85-1.31) | 0.61 | 0.33 | 1.16 (1.00-1.34) | 0.05 | 0.38 | 1.10 (1.01-1.19) | 0.02 | 1.11 (1.04-1.18) | 2.9E-03 | 0.75 |
| rs10022462 | 4 | 89243818 | T | C | 0.18 | 1.29 (1.01-1.66) | 0.04 | 0.28 | 1.07 (0.92-1.25) | 0.35 | 0.29 | 0.94 (0.86-1.03) | 0.19 | 1.00 (0.93-1.07) | 0.94 | 0.04 |
| rs9790517 | 4 | 106084778 | T | C | 0.34 | 1.11 (0.91-1.37) | 0.30 | 0.31 | 1.06 (0.92-1.22) | 0.45 | 0.31 | 1.01 (0.93-1.09) | 0.86 | 1.03 (0.96-1.10) | 0.41 | 0.61 |
| rs77528541 | 4 | 126843504 | G | T | 0.95 | 1.15 (0.74-1.79) | 0.54 | 0.95 | 1.13 (0.82-1.55) | 0.45 | 0.94 | 1.18 (1.00-1.39) | 0.05 | 1.17 (1.01-1.34) | 0.03 | 0.97 |
| rs6828523 | 4 | 175846426 | C | A | 0.78 | 1.04 (0.82-1.32) | 0.74 | 0.87 | 0.78 (0.65-0.95) | 0.01 | 0.81 | 1.06 (0.96-1.16) | 0.27 | 1.00 (0.92-1.08) | 0.99 | 0.02 |
| rs116095464 | 5 | 345109 | C | T | 0.06 | 1.00 (0.65-1.54) | 1.00 | 0.08 | 0.86 (0.66-1.11) | 0.24 | 0.07 | 1.15 (0.99-1.33) | 0.06 | 1.06 (0.94-1.20) | 0.32 | 0.15 |
| rs10069690 | 5 | 1279790 | T | C | 0.18 | 0.86 (0.67-1.11) | 0.26 | 0.20 | 1.09 (0.93-1.29) | 0.29 | 0.21 | 1.10 (1.00-1.21) | 0.06 | 1.07 (0.99-1.16) | 0.08 | 0.22 |
| rs3215401 | 5 | 1296255 | A | AG | 0.78 | 0.99 (0.79-1.25) | 0.95 | 0.76 | 1.15 (0.98-1.36) | 0.09 | 0.75 | 1.00 (0.92-1.10) | 0.93 | 1.03 (0.96-1.11) | 0.42 | 0.33 |
| rs13162653 | 5 | 16187528 | G | T | 0.66 | 0.97 (0.78-1.19) | 0.75 | 0.61 | 0.98 (0.85-1.12) | 0.75 | 0.62 | 1.04 (0.96-1.12) | 0.33 | 1.02 (0.95-1.09) | 0.58 | 0.66 |
| rs2012709 | 5 | 32567732 | T | C | 0.37 | 1.07 (0.88-1.31) | 0.50 | 0.32 | 1.00 (0.86-1.16) | 0.99 | 0.40 | 0.92 (0.85-1.00) | 0.04 | 0.95 (0.89-1.02) | 0.14 | 0.30 |
| rs10941679 | 5 | 44706498 | G | A | 0.38 | 1.08 (0.88-1.32) | 0.47 | 0.31 | 1.22 (1.05-1.41) | 8.3E-03 | 0.32 | 1.09 (1.00-1.18) | 0.04 | 1.11 (1.04-1.19) | 1.7E-03 | 0.40 |
| rs35951924 | 5 | 50195093 | A | AT | 0.81 | 0.92 (0.71-1.19) | 0.54 | 0.84 | 0.87 (0.73-1.05) | 0.16 | 0.76 | 1.02 (0.93-1.12) | 0.61 | 0.99 (0.91-1.07) | 0.72 | 0.28 |
| rs62355902 | 5 | 56053723 | T | A | 0.25 | 1.04 (0.84-1.30) | 0.70 | 0.24 | 1.06 (0.90-1.24) | 0.49 | 0.23 | 1.10 (1.01-1.20) | 0.03 | 1.09 (1.01-1.17) | 0.02 | 0.84 |
| rs10472076 | 5 | 58184061 | C | T | 0.30 | 1.08 (0.88-1.34) | 0.46 | 0.32 | 0.88 (0.76-1.02) | 0.08 | 0.32 | 1.01 (0.93-1.09) | 0.84 | 0.99 (0.92-1.05) | 0.68 | 0.18 |
| rs1353747 | 5 | 58337481 | T | G | 0.89 | 0.94 (0.68-1.28) | 0.68 | 0.90 | 0.92 (0.73-1.15) | 0.46 | 0.91 | 1.01 (0.89-1.14) | 0.92 | 0.98 (0.88-1.09) | 0.69 | 0.76 |
| rs7707921 | 5 | 81538046 | A | T | 0.91 | 0.98 (0.69-1.40) | 0.93 | 0.84 | 1.06 (0.88-1.27) | 0.54 | 0.85 | 1.01 (0.90-1.13) | 0.90 | 1.02 (0.93-1.12) | 0.70 | 0.88 |
| rs10474352 | 5 | 90732225 | C | T | 0.78 | 0.94 (0.74-1.18) | 0.58 | 0.79 | 1.06 (0.90-1.24) | 0.51 | 0.80 | 1.14 (1.04-1.26) | 6.7E-03 | 1.10 (1.01-1.19) | 0.02 | 0.26 |
| rs6882649 | 5 | 111217786 | T | G | 0.43 | 0.82 (0.67-1.00) | 0.05 | 0.51 | 0.91 (0.80-1.05) | 0.20 | 0.50 | 1.08 (1.00-1.16) | 0.07 | 1.01 (0.95-1.08) | 0.78 | 0.01 |
| rs6596100 | 5 | 132407058 | C | T | 0.91 | 1.20 (0.84-1.70) | 0.31 | 0.87 | 1.09 (0.89-1.33) | 0.41 | 0.86 | 1.10 (0.98-1.24) | 0.10 | 1.11 (1.00-1.22) | 0.04 | 0.89 |
| rs1432679 | 5 | 158244083 | C | T | 0.55 | 0.83 (0.68-1.01) | 0.07 | 0.64 | 1.09 (0.95-1.26) | 0.22 | 0.53 | 1.08 (1.00-1.16) | 0.05 | 1.05 (0.99-1.12) | 0.10 | 0.05 |
| rs4562056 | 5 | 169591487 | T | G | 0.48 | 1.02 (0.84-1.24) | 0.83 | 0.38 | 1.06 (0.92-1.21) | 0.43 | 0.44 | 1.02 (0.95-1.10) | 0.53 | 1.03 (0.97-1.10) | 0.35 | 0.92 |
| rs11242675 | 6 | 1318878 | T | C | 0.48 | 1.04 (0.86-1.26) | 0.71 | 0.52 | 0.95 (0.83-1.09) | 0.47 | 0.55 | 1.05 (0.97-1.13) | 0.21 | 1.03 (0.96-1.09) | 0.41 | 0.46 |
| rs204247 | 6 | 13722523 | G | A | 0.41 | 1.10 (0.91-1.33) | 0.34 | 0.51 | 0.91 (0.79-1.04) | 0.17 | 0.42 | 1.06 (0.99-1.15) | 0.11 | 1.03 (0.97-1.10) | 0.32 | 0.11 |
| rs3819405 | 6 | 16399557 | C | T | 0.50 | 0.98 (0.81-1.19) | 0.86 | 0.57 | 0.96 (0.84-1.10) | 0.57 | 0.53 | 1.06 (0.98-1.14) | 0.15 | 1.03 (0.97-1.10) | 0.37 | 0.43 |
| rs2223621 | 6 | 20621238 | T | C | 0.35 | 1.05 (0.86-1.28) | 0.62 | 0.29 | 1.12 (0.97-1.30) | 0.12 | 0.34 | 0.99 (0.92-1.07) | 0.85 | 1.02 (0.96-1.09) | 0.49 | 0.34 |
| rs71557345 | 6 | 26680698 | G | A | 0.98 | 0.81 (0.39-1.67) | 0.56 | 0.98 | 0.90 (0.53-1.54) | 0.70 | 0.97 | 1.00 (0.76-1.30) | 0.99 | 0.96 (0.76-1.20) | 0.72 | 0.84 |
| rs9257408 | 6 | 28926220 | C | G | 0.54 | 1.10 (0.91-1.33) | 0.32 | 0.58 | 0.93 (0.81-1.07) | 0.31 | 0.63 | 1.03 (0.94-1.13) | 0.50 | 1.01 (0.95-1.09) | 0.71 | 0.31 |
| rs12207986 | 6 | 81094287 | A | G | 0.22 | 0.77 (0.60-0.98) | 0.04 | 0.30 | 1.03 (0.89-1.19) | 0.69 | 0.32 | 0.92 (0.85-1.01) | 0.08 | 0.93 (0.87-1.00) | 0.06 | 0.12 |
| rs17529111 | 6 | 82128386 | C | T | 0.26 | 1.01 (0.82-1.25) | 0.89 | 0.12 | 1.16 (0.95-1.43) | 0.15 | 0.24 | 1.13 (1.03-1.23) | 6.8E-03 | 1.12 (1.04-1.20) | 3.7E-03 | 0.62 |
| rs6569648 | 6 | 130349119 | T | C | 0.93 | 1.33 (0.92-1.94) | 0.13 | 0.86 | 1.06 (0.87-1.30) | 0.56 | 0.88 | 0.91 (0.81-1.03) | 0.14 | 0.97 (0.88-1.08) | 0.61 | 0.10 |
| rs9485372 | 6 | 149608874 | G | A | 0.85 | 1.06 (0.82-1.37) | 0.67 | 0.83 | 0.93 (0.78-1.11) | 0.43 | 0.82 | 1.05 (0.96-1.16) | 0.29 | 1.03 (0.95-1.12) | 0.51 | 0.47 |
| rs3757322 | 6 | 151942194 | G | T | 0.18 | 1.01 (0.78-1.30) | 0.95 | 0.23 | 1.17 (1.00-1.37) | 0.06 | 0.25 | 1.14 (1.04-1.25) | 3.9E-03 | 1.13 (1.05-1.22) | 8.9E-04 | 0.62 |
| rs9397437 | 6 | 151952332 | A | G | 0.03 | 1.52 (0.90-2.57) | 0.12 | 0.04 | 1.40 (1.02-1.92) | 0.04 | 0.05 | 1.17 (0.99-1.38) | 0.06 | 1.24 (1.07-1.42) | 3.3E-03 | 0.46 |
| rs140068132 | 6 | 151954834 | A | G | 0.86 | 1.02 (0.77-1.35) | 0.90 | 0.89 | 1.45 (1.15-1.83) | 1.9E-03 | 0.90 | 1.70 (1.49-1.95) | 5.6E-15 | 1.53 (1.37-1.70) | 9.4E-15 | 4.80E-03 |
| rs851984 | 6 | 152023191 | A | G | 0.38 | 0.89 (0.73-1.09) | 0.27 | 0.33 | 1.12 (0.97-1.30) | 0.11 | 0.35 | 1.25 (1.15-1.35) | 3.6E-08 | 1.18 (1.10-1.26) | 8.4E-07 | 0.01 |
| rs3778609 | 6 | 152133187 | C | T | 0.74 | 1.08 (0.86-1.35) | 0.52 | 0.81 | 1.12 (0.94-1.34) | 0.21 | 0.80 | 1.34 (1.21-1.47) | 2.3E-09 | 1.26 (1.16-1.36) | 1.1E-08 | 0.08 |
| rs2747652 | 6 | 152437016 | C | T | 0.53 | 0.97 (0.80-1.17) | 0.72 | 0.50 | 1.02 (0.89-1.16) | 0.79 | 0.51 | 1.05 (0.97-1.13) | 0.24 | 1.03 (0.97-1.10) | 0.33 | 0.73 |
| rs7971 | 7 | 21940960 | A | G | 0.76 | 1.08 (0.87-1.35) | 0.49 | 0.68 | 0.95 (0.82-1.09) | 0.45 | 0.71 | 0.97 (0.90-1.06) | 0.53 | 0.98 (0.91-1.05) | 0.51 | 0.61 |
| rs17156577 | 7 | 28356889 | C | T | 0.20 | 0.81 (0.63-1.03) | 0.08 | 0.15 | 0.85 (0.70-1.03) | 0.10 | 0.17 | 0.96 (0.87-1.05) | 0.37 | 0.92 (0.84-1.00) | 0.04 | 0.31 |

| | | | | | | | | | | | | | | | | |
|---|---|---|---|---|---|---|---|---|---|---|---|---|---|---|---|---|
| rs6964587 | 7 | 91630620 | T | G | 0.30 | 1.12 (0.90-1.39) | 0.30 | 0.37 | 0.96 (0.84-1.11) | 0.60 | 0.35 | 1.01 (0.93-1.09) | 0.77 | 1.01 (0.95-1.08) | 0.75 | 0.51 |
| rs17268829 | 7 | 94113799 | C | T | 0.35 | 1.10 (0.90-1.34) | 0.35 | 0.28 | 1.03 (0.89-1.20) | 0.67 | 0.33 | 1.00 (0.92-1.08) | 0.92 | 1.01 (0.95-1.08) | 0.68 | 0.64 |
| rs71559437 | 7 | 101552440 | G | A | 0.94 | 0.78 (0.51-1.20) | 0.26 | 0.91 | 1.03 (0.82-1.31) | 0.79 | 0.92 | 0.99 (0.87-1.14) | 0.94 | 0.99 (0.88-1.11) | 0.82 | 0.52 |
| rs4593472 | 7 | 130667121 | C | T | 0.85 | 0.97 (0.75-1.27) | 0.85 | 0.85 | 1.11 (0.92-1.34) | 0.27 | 0.79 | 0.98 (0.89-1.08) | 0.68 | 1.00 (0.92-1.09) | 0.94 | 0.49 |
| rs11977670 | 7 | 139942304 | A | G | 0.63 | 0.97 (0.79-1.18) | 0.75 | 0.54 | 1.01 (0.88-1.15) | 0.94 | 0.55 | 1.01 (0.93-1.09) | 0.83 | 1.00 (0.94-1.07) | 0.91 | 0.93 |
| rs720475 | 7 | 144074929 | G | A | 0.88 | 1.00 (0.75-1.33) | 0.99 | 0.88 | 0.92 (0.75-1.12) | 0.40 | 0.83 | 1.05 (0.95-1.17) | 0.34 | 1.02 (0.93-1.12) | 0.67 | 0.49 |
| rs66823261 | 8 | 170692 | C | T | 0.20 | 0.92 (0.72-1.17) | 0.50 | 0.22 | 0.97 (0.82-1.14) | 0.71 | 0.22 | 1.19 (1.09-1.30) | 1.7E-04 | 1.11 (1.03-1.20) | 5.7E-03 | 0.03 |
| rs9693444 | 8 | 29509616 | A | C | 0.32 | 0.96 (0.78-1.18) | 0.72 | 0.28 | 1.01 (0.87-1.18) | 0.89 | 0.33 | 0.97 (0.90-1.05) | 0.48 | 0.98 (0.91-1.05) | 0.52 | 0.89 |
| rs13365225 | 8 | 36858483 | A | G | 0.83 | 1.09 (0.84-1.42) | 0.51 | 0.76 | 1.10 (0.94-1.29) | 0.25 | 0.81 | 1.07 (0.97-1.18) | 0.18 | 1.08 (1.00-1.17) | 0.06 | 0.95 |
| rs6472903 | 8 | 76230301 | T | G | 0.94 | 0.95 (0.64-1.40) | 0.79 | 0.94 | 0.90 (0.67-1.19) | 0.45 | 0.91 | 1.06 (0.92-1.22) | 0.41 | 1.02 (0.91-1.15) | 0.75 | 0.54 |
| rs2943559 | 8 | 76417937 | G | A | 0.05 | 1.18 (0.77-1.79) | 0.45 | 0.08 | 0.96 (0.74-1.23) | 0.73 | 0.07 | 0.97 (0.84-1.12) | 0.70 | 0.98 (0.87-1.11) | 0.79 | 0.68 |
| rs514192 | 8 | 102478959 | A | T | 0.41 | 1.04 (0.86-1.26) | 0.69 | 0.41 | 1.02 (0.89-1.17) | 0.81 | 0.39 | 1.05 (0.97-1.13) | 0.25 | 1.04 (0.98-1.11) | 0.24 | 0.94 |
| rs12546444 | 8 | 106358620 | A | T | 0.90 | 1.05 (0.77-1.45) | 0.75 | 0.93 | 0.97 (0.75-1.27) | 0.85 | 0.90 | 1.09 (0.96-1.23) | 0.20 | 1.06 (0.96-1.18) | 0.26 | 0.77 |
| rs13267382 | 8 | 117209548 | A | G | 0.69 | 0.93 (0.75-1.16) | 0.53 | 0.62 | 1.04 (0.91-1.20) | 0.55 | 0.61 | 1.07 (0.99-1.16) | 0.11 | 1.05 (0.98-1.12) | 0.15 | 0.52 |
| rs58847541 | 8 | 124610166 | A | G | 0.07 | 1.00 (0.68-1.47) | 0.98 | 0.09 | 0.99 (0.78-1.25) | 0.92 | 0.08 | 1.00 (0.87-1.16) | 0.95 | 1.00 (0.89-1.13) | 1.00 | 0.99 |
| rs17350191 | 8 | 124757661 | T | C | 0.29 | 1.10 (0.89-1.35) | 0.39 | 0.27 | 1.01 (0.87-1.18) | 0.89 | 0.31 | 1.08 (1.00-1.17) | 0.06 | 1.07 (1.00-1.14) | 0.06 | 0.73 |
| rs13281615 | 8 | 128355618 | G | A | 0.67 | 0.91 (0.74-1.13) | 0.39 | 0.54 | 1.12 (0.98-1.28) | 0.10 | 0.59 | 1.12 (1.04-1.21) | 3.8E-03 | 1.10 (1.03-1.17) | 3.5E-03 | 0.19 |
| rs11780156 | 8 | 129194641 | T | C | 0.18 | 0.94 (0.73-1.22) | 0.64 | 0.15 | 1.31 (1.10-1.57) | 2.9E-03 | 0.17 | 1.06 (0.96-1.17) | 0.24 | 1.10 (1.01-1.19) | 0.03 | 0.06 |
| rs1011970 | 9 | 22062134 | T | G | 0.43 | 1.15 (0.95-1.41) | 0.16 | 0.34 | 1.13 (0.97-1.30) | 0.11 | 0.34 | 1.06 (0.98-1.14) | 0.17 | 1.08 (1.01-1.15) | 0.02 | 0.60 |
| rs10759243 | 9 | 110306115 | A | C | 0.49 | 1.03 (0.85-1.25) | 0.77 | 0.41 | 0.89 (0.77-1.03) | 0.11 | 0.41 | 1.12 (1.04-1.21) | 2.5E-03 | 1.06 (1.00-1.13) | 0.06 | 0.02 |
| rs10816625 | 9 | 110837073 | G | A | 0.19 | 1.14 (0.90-1.44) | 0.29 | 0.13 | 1.01 (0.83-1.23) | 0.93 | 0.15 | 1.10 (1.00-1.21) | 0.06 | 1.09 (1.00-1.18) | 0.05 | 0.70 |
| rs13294895 | 9 | 110837176 | T | C | 0.11 | 0.78 (0.55-1.09) | 0.14 | 0.08 | 0.95 (0.74-1.22) | 0.70 | 0.12 | 1.10 (0.97-1.23) | 0.13 | 1.04 (0.94-1.15) | 0.48 | 0.13 |
| rs676256 | 9 | 110895353 | T | C | 0.73 | 1.09 (0.88-1.36) | 0.42 | 0.68 | 1.01 (0.87-1.16) | 0.94 | 0.68 | 1.14 (1.05-1.24) | 1.5E-03 | 1.11 (1.03-1.18) | 3.6E-03 | 0.31 |
| rs1895062 | 9 | 119313486 | A | G | 0.72 | 0.98 (0.79-1.22) | 0.88 | 0.66 | 1.11 (0.96-1.28) | 0.17 | 0.67 | 1.08 (0.99-1.17) | 0.08 | 1.07 (1.00-1.15) | 0.04 | 0.66 |
| rs10760444 | 9 | 129396434 | G | A | 0.49 | 0.87 (0.72-1.06) | 0.18 | 0.57 | 1.04 (0.91-1.20) | 0.55 | 0.48 | 1.02 (0.94-1.10) | 0.65 | 1.01 (0.95-1.07) | 0.82 | 0.32 |
| rs8176636 | 9 | 136151579 | T | TGGTGCAGGCGCAGGAAAAAATTGTGGCAATTCCTCA | 0.11 | 1.09 (0.80-1.49) | 0.56 | 0.15 | 0.80 (0.66-0.97) | 0.03 | 0.15 | 1.11 (0.99-1.23) | 0.06 | 1.03 (0.94-1.13) | 0.48 | 0.02 |
| rs2380205 | 10 | 5886734 | C | T | 0.75 | 1.09 (0.87-1.36) | 0.47 | 0.69 | 1.03 (0.89-1.19) | 0.73 | 0.67 | 0.98 (0.90-1.07) | 0.65 | 1.00 (0.93-1.07) | 1.00 | 0.66 |
| rs67958007 | 10 | 9088113 | T | TG | 0.20 | 0.99 (0.78-1.26) | 0.93 | 0.13 | 0.99 (0.81-1.21) | 0.90 | 0.17 | 1.02 (0.92-1.12) | 0.75 | 1.01 (0.93-1.09) | 0.85 | 0.96 |
| rs7072776 | 10 | 22032942 | A | G | 0.34 | 1.05 (0.85-1.29) | 0.67 | 0.35 | 1.15 (1.00-1.32) | 0.05 | 0.33 | 1.05 (0.97-1.14) | 0.20 | 1.07 (1.00-1.15) | 0.04 | 0.56 |
| rs11814448 | 10 | 22315843 | C | A | 0.04 | 1.49 (0.89-2.50) | 0.13 | 0.06 | 1.12 (0.84-1.50) | 0.43 | 0.06 | 1.16 (0.99-1.37) | 0.07 | 1.18 (1.02-1.35) | 0.02 | 0.63 |
| rs10995201 | 10 | 64299890 | A | G | 0.93 | 0.83 (0.56-1.22) | 0.34 | 0.92 | 0.92 (0.72-1.18) | 0.51 | 0.90 | 1.05 (0.92-1.21) | 0.49 | 1.00 (0.89-1.12) | 0.99 | 0.40 |
| rs704010 | 10 | 80841148 | T | C | 0.41 | 1.16 (0.95-1.41) | 0.15 | 0.44 | 0.99 (0.86-1.13) | 0.86 | 0.41 | 1.06 (0.98-1.15) | 0.13 | 1.05 (0.99-1.12) | 0.10 | 0.42 |
| rs140936696 | 10 | 95292187 | CAA | C | 0.10 | 1.14 (0.83-1.58) | 0.43 | 0.12 | 0.97 (0.79-1.20) | 0.80 | 0.13 | 0.98 (0.88-1.10) | 0.79 | 1.00 (0.90-1.09) | 0.92 | 0.69 |
| rs7904519 | 10 | 114773927 | G | A | 0.27 | 0.94 (0.75-1.19) | 0.63 | 0.30 | 0.93 (0.80-1.08) | 0.33 | 0.35 | 1.08 (0.99-1.17) | 0.08 | 1.03 (0.96-1.10) | 0.38 | 0.17 |
| rs11199914 | 10 | 123093901 | C | T | 0.48 | 0.84 (0.69-1.02) | 0.08 | 0.59 | 1.02 (0.89-1.17) | 0.77 | 0.56 | 0.99 (0.92-1.07) | 0.88 | 0.98 (0.92-1.05) | 0.59 | 0.24 |
| rs35054928 | 10 | 123340431 | GC | G | 0.39 | 1.29 (1.06-1.57) | 0.01 | 0.39 | 1.25 (1.09-1.43) | 1.7E-03 | 0.42 | 1.19 (1.10-1.28) | 7.2E-06 | 1.21 (1.14-1.29) | 2.3E-09 | 0.67 |
| rs45631563 | 10 | 123349324 | A | T | 0.98 | 0.76 (0.35-1.65) | 0.49 | 0.97 | 1.40 (0.89-2.20) | 0.15 | 0.97 | 1.17 (0.93-1.47) | 0.18 | 1.18 (0.96-1.43) | 0.11 | 0.41 |
| rs6597981 | 11 | 803017 | G | A | 0.66 | 0.86 (0.69-1.06) | 0.17 | 0.58 | 0.98 (0.86-1.13) | 0.81 | 0.59 | 1.05 (0.97-1.13) | 0.27 | 1.01 (0.95-1.08) | 0.70 | 0.22 |
| rs3817198 | 11 | 1909006 | C | T | 0.18 | 0.89 (0.69-1.14) | 0.35 | 0.19 | 1.17 (0.99-1.39) | 0.07 | 0.21 | 1.13 (1.03-1.24) | 0.01 | 1.11 (1.03-1.20) | 8.6E-03 | 0.17 |
| rs3903072 | 11 | 65583066 | G | T | 0.77 | 1.15 (0.91-1.45) | 0.26 | 0.71 | 0.99 (0.85-1.15) | 0.91 | 0.68 | 1.01 (0.93-1.10) | 0.87 | 1.02 (0.95-1.09) | 0.67 | 0.56 |

| | | | | | | | | | | | | | | | | |
|---|---|---|---|---|---|---|---|---|---|---|---|---|---|---|---|---|
| rs75915166 | 11 | 69379161 | A | C | 0.01 | 1.78 (0.83-3.82) | 0.14 | 0.01 | 1.51 (0.83-2.76) | 0.18 | 0.03 | 1.30 (1.04-1.63) | 0.02 | 1.35 (1.10-1.66) | 3.5E-03 | 0.69 |
| rs11374964 | 11 | 108345515 | G | GA | 0.42 | 0.94 (0.77-1.15) | 0.56 | 0.43 | 1.03 (0.90-1.18) | 0.64 | 0.46 | 1.04 (0.97-1.12) | 0.29 | 1.03 (0.97-1.10) | 0.36 | 0.65 |
| rs74911261 | 11 | 108357137 | G | A | 0.99 | 0.94 (0.35-2.56) | 0.91 | 0.98 | 1.10 (0.69-1.76) | 0.70 | 0.99 | 0.98 (0.68-1.40) | 0.90 | 1.01 (0.77-1.33) | 0.92 | 0.92 |
| rs11820646 | 11 | 129461171 | C | T | 0.48 | 1.02 (0.84-1.24) | 0.84 | 0.50 | 1.03 (0.90-1.18) | 0.63 | 0.54 | 1.07 (0.99-1.15) | 0.09 | 1.05 (0.99-1.12) | 0.09 | 0.86 |
| rs12422552 | 12 | 14413931 | C | G | 0.18 | 1.21 (0.94-1.55) | 0.14 | 0.21 | 0.96 (0.81-1.13) | 0.60 | 0.22 | 1.07 (0.98-1.18) | 0.13 | 1.06 (0.98-1.14) | 0.14 | 0.27 |
| rs7297051 | 12 | 28174817 | C | T | 0.59 | 0.68 (0.55-0.84) | 2.8E-04 | 0.75 | 1.10 (0.94-1.29) | 0.24 | 0.65 | 1.15 (1.06-1.24) | 6.2E-04 | 1.08 (1.01-1.15) | 0.03 | 2.33E-05 |
| rs202049448 | 12 | 85009437 | T | C | 0.66 | 1.13 (0.93-1.38) | 0.22 | 0.65 | 0.96 (0.83-1.10) | 0.54 | 0.64 | 1.00 (0.92-1.08) | 1.00 | 1.00 (0.94-1.07) | 0.91 | 0.39 |
| rs17356907 | 12 | 96027759 | A | G | 0.59 | 1.04 (0.85-1.26) | 0.72 | 0.61 | 1.07 (0.93-1.23) | 0.34 | 0.64 | 1.06 (0.99-1.15) | 0.11 | 1.06 (1.00-1.13) | 0.06 | 0.97 |
| rs1292011 | 12 | 115836522 | A | G | 0.65 | 0.89 (0.73-1.09) | 0.25 | 0.61 | 1.14 (0.99-1.32) | 0.06 | 0.63 | 1.00 (0.92-1.08) | 0.95 | 1.01 (0.95-1.08) | 0.65 | 0.10 |
| rs206966 | 12 | 120832146 | T | C | 0.07 | 1.12 (0.75-1.67) | 0.59 | 0.10 | 1.03 (0.81-1.29) | 0.83 | 0.09 | 1.03 (0.90-1.18) | 0.68 | 1.04 (0.92-1.16) | 0.55 | 0.93 |
| rs11571833 | 13 | 32972626 | T | A | 0.00 | 0.69 (0.04-11.14) | 0.79 | 0.00 | 1.00 (0.37-2.70) | 1.00 | 0.00 | 1.01 (0.54-1.89) | 0.98 | 0.99 (0.59-1.67) | 0.98 | 0.97 |
| rs6562760 | 13 | 73957681 | G | A | 0.89 | 1.18 (0.86-1.62) | 0.32 | 0.83 | 1.18 (0.99-1.42) | 0.07 | 0.82 | 1.04 (0.94-1.16) | 0.46 | 1.08 (0.99-1.18) | 0.08 | 0.43 |
| rs2236007 | 14 | 37132769 | G | A | 0.89 | 0.60 (0.42-0.86) | 5.3E-03 | 0.87 | 1.09 (0.89-1.34) | 0.41 | 0.87 | 1.14 (1.01-1.29) | 0.04 | 1.07 (0.97-1.19) | 0.17 | 4.10E-03 |
| rs2588809 | 14 | 68660428 | T | C | 0.13 | 1.11 (0.83-1.47) | 0.48 | 0.17 | 1.10 (0.92-1.32) | 0.29 | 0.17 | 1.05 (0.94-1.16) | 0.38 | 1.07 (0.98-1.16) | 0.15 | 0.86 |
| rs999737 | 14 | 69034682 | C | T | 0.86 | 1.09 (0.83-1.43) | 0.53 | 0.85 | 1.04 (0.85-1.26) | 0.70 | 0.83 | 0.98 (0.89-1.09) | 0.76 | 1.00 (0.92-1.09) | 0.91 | 0.74 |
| rs941764 | 14 | 91841069 | G | A | 0.51 | 0.92 (0.76-1.11) | 0.39 | 0.41 | 1.01 (0.88-1.16) | 0.91 | 0.46 | 1.06 (0.98-1.14) | 0.14 | 1.03 (0.97-1.10) | 0.33 | 0.38 |
| rs11627032 | 14 | 93104072 | T | C | 0.71 | 0.90 (0.73-1.12) | 0.36 | 0.70 | 1.05 (0.91-1.22) | 0.50 | 0.73 | 0.99 (0.91-1.08) | 0.88 | 1.00 (0.93-1.07) | 0.91 | 0.52 |
| rs10623258 | 14 | 105212261 | CTT | C | 0.71 | 1.01 (0.81-1.27) | 0.89 | 0.63 | 1.07 (0.93-1.23) | 0.35 | 0.62 | 1.00 (0.92-1.08) | 0.99 | 1.02 (0.95-1.08) | 0.64 | 0.71 |
| rs2290203 | 15 | 91512067 | G | A | 0.63 | 0.89 (0.72-1.10) | 0.29 | 0.73 | 1.03 (0.89-1.20) | 0.70 | 0.68 | 1.17 (1.08-1.27) | 1.4E-04 | 1.11 (1.04-1.19) | 2.6E-03 | 0.03 |
| rs11076805 | 16 | 4106788 | C | A | 0.91 | 1.18 (0.84-1.64) | 0.34 | 0.85 | 1.25 (1.02-1.52) | 0.03 | 0.85 | 1.13 (1.01-1.26) | 0.04 | 1.16 (1.05-1.27) | 2.5E-03 | 0.68 |
| rs4784227 | 16 | 52599188 | T | C | 0.31 | 1.53 (1.24-1.89) | 7.0E-05 | 0.35 | 1.21 (1.05-1.39) | 8.2E-03 | 0.29 | 1.25 (1.15-1.36) | 4.5E-08 | 1.27 (1.19-1.35) | 2.4E-12 | 0.16 |
| rs17817449 | 16 | 53813367 | T | G | 0.81 | 0.97 (0.76-1.25) | 0.82 | 0.77 | 0.95 (0.81-1.11) | 0.52 | 0.73 | 1.04 (0.96-1.14) | 0.34 | 1.02 (0.94-1.09) | 0.67 | 0.55 |
| rs11075995 | 16 | 53855291 | A | T | 0.41 | 0.91 (0.75-1.10) | 0.33 | 0.27 | 1.22 (1.05-1.41) | 9.4E-03 | 0.32 | 1.10 (1.02-1.19) | 0.02 | 1.10 (1.03-1.17) | 5.3E-03 | 0.06 |
| rs28539243 | 16 | 54682064 | A | G | 0.58 | 0.92 (0.76-1.11) | 0.39 | 0.59 | 0.92 (0.80-1.06) | 0.24 | 0.57 | 0.99 (0.91-1.06) | 0.73 | 0.97 (0.91-1.03) | 0.27 | 0.61 |
| rs2432539 | 16 | 56420987 | A | G | 0.37 | 0.96 (0.79-1.17) | 0.69 | 0.45 | 1.07 (0.94-1.23) | 0.32 | 0.38 | 1.02 (0.95-1.10) | 0.55 | 1.03 (0.96-1.09) | 0.41 | 0.66 |
| rs13329835 | 16 | 80650805 | G | A | 0.12 | 1.41 (1.05-1.91) | 0.02 | 0.19 | 1.08 (0.91-1.27) | 0.39 | 0.19 | 1.00 (0.90-1.10) | 0.97 | 1.04 (0.96-1.13) | 0.30 | 0.09 |
| rs4496150 | 16 | 87085237 | C | A | 0.59 | 0.94 (0.77-1.15) | 0.57 | 0.65 | 1.15 (1.00-1.33) | 0.05 | 0.65 | 0.96 (0.89-1.04) | 0.30 | 1.00 (0.93-1.06) | 0.90 | 0.07 |
| rs146699004 | 17 | 29230520 | GGT | G | 0.87 | 0.84 (0.62-1.13) | 0.25 | 0.81 | 0.99 (0.84-1.18) | 0.93 | 0.82 | 1.01 (0.91-1.11) | 0.92 | 0.99 (0.91-1.08) | 0.78 | 0.53 |
| rs72826962 | 17 | 40836389 | T | C | 0.01 | 0.59 (0.10-3.55) | 0.57 | 0.01 | 0.65 (0.24-1.75) | 0.39 | 0.01 | 1.35 (0.87-2.10) | 0.18 | 1.16 (0.78-1.72) | 0.45 | 0.31 |
| rs2532263 | 17 | 44252468 | G | A | 0.90 | 1.14 (0.82-1.59) | 0.44 | 0.86 | 1.04 (0.85-1.27) | 0.68 | 0.88 | 0.99 (0.86-1.13) | 0.85 | 1.02 (0.91-1.13) | 0.76 | 0.71 |
| rs2787486 | 17 | 53209774 | A | C | 0.81 | 1.19 (0.92-1.54) | 0.18 | 0.82 | 1.06 (0.89-1.27) | 0.51 | 0.76 | 1.05 (0.96-1.15) | 0.30 | 1.06 (0.98-1.15) | 0.12 | 0.65 |
| rs745570 | 17 | 77781725 | A | G | 0.27 | 0.88 (0.70-1.11) | 0.28 | 0.33 | 1.13 (0.98-1.30) | 0.10 | 0.35 | 1.01 (0.93-1.10) | 0.77 | 1.02 (0.96-1.10) | 0.48 | 0.17 |
| rs527616 | 18 | 24337424 | G | C | 0.84 | 0.81 (0.62-1.06) | 0.13 | 0.76 | 1.01 (0.87-1.19) | 0.86 | 0.77 | 1.09 (0.99-1.20) | 0.07 | 1.05 (0.97-1.13) | 0.26 | 0.12 |
| rs1436904 | 18 | 24570667 | T | G | 0.55 | 1.11 (0.92-1.35) | 0.27 | 0.62 | 0.96 (0.83-1.10) | 0.52 | 0.56 | 1.08 (1.00-1.17) | 0.04 | 1.06 (0.99-1.13) | 0.07 | 0.25 |
| rs36194942 | 18 | 25401204 | A | AT | 0.76 | 0.96 (0.77-1.21) | 0.75 | 0.72 | 1.11 (0.95-1.29) | 0.20 | 0.71 | 1.15 (1.06-1.25) | 1.2E-03 | 1.12 (1.04-1.20) | 1.5E-03 | 0.35 |
| rs117618124 | 18 | 29977689 | T | C | 1.00 | 0.72 (0.14-3.71) | 0.69 | 0.99 | 1.39 (0.63-3.07) | 0.42 | 0.98 | 1.49 (1.04-2.13) | 0.03 | 1.43 (1.04-1.97) | 0.03 | 0.70 |
| rs6507583 | 18 | 42399590 | A | G | 0.92 | 0.92 (0.64-1.32) | 0.65 | 0.92 | 1.17 (0.90-1.51) | 0.25 | 0.91 | 1.05 (0.92-1.20) | 0.49 | 1.06 (0.94-1.18) | 0.34 | 0.57 |
| rs322144 | 19 | 11423703 | C | G | 0.31 | 0.84 (0.68-1.04) | 0.11 | 0.42 | 0.87 (0.76-1.00) | 0.05 | 0.40 | 0.95 (0.88-1.03) | 0.23 | 0.92 (0.86-0.98) | 0.02 | 0.34 |
| rs78269692 | 19 | 13158277 | C | T | 0.01 | 0.74 (0.28-1.96) | 0.54 | 0.01 | 1.74 (0.89-3.40) | 0.10 | 0.02 | 0.85 (0.61-1.21) | 0.37 | 0.97 (0.72-1.29) | 0.82 | 0.15 |
| rs2594714 | 19 | 13954571 | G | A | 0.70 | 1.14 (0.92-1.41) | 0.23 | 0.72 | 0.99 (0.86-1.15) | 0.94 | 0.72 | 1.00 (0.92-1.08) | 0.93 | 1.01 (0.94-1.08) | 0.79 | 0.51 |
| rs67397200 | 19 | 17401404 | G | C | 0.10 | 1.08 (0.79-1.49) | 0.62 | 0.16 | 0.97 (0.80-1.17) | 0.72 | 0.18 | 0.94 (0.85-1.05) | 0.26 | 0.96 (0.88-1.04) | 0.33 | 0.71 |
| rs4808801 | 19 | 18571141 | A | G | 0.60 | 0.96 (0.78-1.17) | 0.66 | 0.68 | 1.03 (0.89-1.19) | 0.70 | 0.62 | 1.12 (1.03-1.21) | 5.3E-03 | 1.08 (1.01-1.15) | 0.02 | 0.28 |
| rs2965183 | 19 | 19545696 | A | G | 0.49 | 0.90 (0.74-1.09) | 0.28 | 0.43 | 0.98 (0.86-1.13) | 0.82 | 0.42 | 1.04 (0.97-1.12) | 0.27 | 1.02 (0.95-1.08) | 0.64 | 0.33 |
| rs113701136 | 19 | 30277729 | T | C | 0.27 | 0.76 (0.60-0.96) | 0.02 | 0.32 | 1.00 (0.87-1.15) | 0.99 | 0.27 | 0.96 (0.88-1.04) | 0.30 | 0.95 (0.88-1.02) | 0.12 | 0.13 |
| rs3760982 | 19 | 44286513 | A | G | 0.26 | 1.09 (0.87-1.37) | 0.44 | 0.29 | 1.07 (0.92-1.25) | 0.36 | 0.33 | 1.05 (0.97-1.14) | 0.25 | 1.06 (0.99-1.13) | 0.11 | 0.93 |

| | | | | | | | | | | | | | | | | |
|---|---|---|---|---|---|---|---|---|---|---|---|---|---|---|---|---|
| rs71338792 | 19 | 46183031 | AT | A | 0.10 | 1.25 (0.92-1.70) | 0.15 | 0.12 | 1.13 (0.92-1.39) | 0.24 | 0.13 | 1.05 (0.93-1.18) | 0.42 | 1.09 (0.99-1.20) | 0.09 | 0.52 |
| rs16991615 | 20 | 5948227 | A | G | 0.05 | 0.77 (0.49-1.22) | 0.27 | 0.06 | 0.99 (0.74-1.33) | 0.96 | 0.05 | 1.05 (0.88-1.25) | 0.60 | 1.00 (0.87-1.16) | 0.96 | 0.47 |
| rs2284378 | 20 | 32588095 | T | C | 0.39 | 0.79 (0.65-0.98) | 0.03 | 0.31 | 0.96 (0.83-1.11) | 0.59 | 0.31 | 1.09 (1.00-1.18) | 0.04 | 1.03 (0.96-1.10) | 0.45 | 0.01 |
| rs6122906 | 20 | 48945911 | G | A | 0.40 | 0.87 (0.71-1.06) | 0.18 | 0.33 | 1.06 (0.92-1.23) | 0.39 | 0.30 | 1.02 (0.94-1.11) | 0.58 | 1.01 (0.95-1.08) | 0.69 | 0.26 |
| rs2823093 | 21 | 16520832 | G | A | 0.70 | 1.06 (0.86-1.31) | 0.57 | 0.74 | 1.00 (0.86-1.17) | 0.98 | 0.70 | 1.00 (0.93-1.09) | 0.94 | 1.01 (0.94-1.08) | 0.79 | 0.88 |
| rs132390 | 22 | 29621477 | C | T | 0.00 | 2.30 (0.45-11.83) | 0.32 | 0.01 | 1.46 (0.73-2.91) | 0.28 | 0.01 | 0.55 (0.36-0.86) | 8.4E-03 | 0.77 (0.54-1.11) | 0.16 | 0.03 |
| rs738321 | 22 | 38568833 | C | G | 0.55 | 0.92 (0.76-1.11) | 0.39 | 0.53 | 1.24 (1.08-1.42) | 2.5E-03 | 0.57 | 1.01 (0.94-1.09) | 0.71 | 1.05 (0.98-1.11) | 0.16 | 0.02 |
| chr22:39359355 | 22 | 39359355 | <CN0> | C | 0.26 | 1.00 (0.81-1.25) | 0.99 | 0.19 | 1.06 (0.89-1.25) | 0.52 | 0.18 | 1.09 (0.99-1.20) | 0.08 | 1.07 (0.99-1.16) | 0.09 | 0.79 |
| rs6001930 | 22 | 40876234 | C | T | 0.08 | 1.17 (0.83-1.66) | 0.36 | 0.10 | 1.21 (0.96-1.51) | 0.10 | 0.09 | 1.06 (0.93-1.21) | 0.35 | 1.10 (0.99-1.23) | 0.07 | 0.59 |
| rs73161324 | 22 | 42038786 | T | C | 0.01 | 0.55 (0.16-1.92) | 0.35 | 0.02 | 0.71 (0.41-1.21) | 0.20 | 0.02 | 0.77 (0.56-1.04) | 0.09 | 0.74 (0.57-0.96) | 0.02 | 0.86 |
| rs28512361 | 22 | 46283297 | A | G | 0.06 | 0.98 (0.65-1.48) | 0.92 | 0.07 | 1.01 (0.78-1.32) | 0.92 | 0.08 | 0.85 (0.73-0.99) | 0.04 | 0.90 (0.79-1.02) | 0.09 | 0.48 |

* rs72749841 not included in meta-analysis due to imputation quality
† SFBCS/NC-BCFR, RPGEH, MEC, CAMA, PEGEN-BC

**Table S3.** Association of PRS constructed using different imputation $r^2$ cutoffs with breast cancer risk

| Imputation $r^2$ cutoff | Number of SNPs | OR per SD (95% CI)* |
|---|---|---|
| $r^2 > 0$ | 180 | 1.57 (1.49 to 1.65) |
| $r^2 > 0.5$ | 166 | 1.57 (1.50 to 1.66) |
| $r^2 > 0.8$ | 134 | 1.53 (1.45 to 1.61) |

* Odds ratio from multivariable logistic regression of PRS adjusted for study and genetic ancestry among participants of SFBCS/NC-BCFR, RPGEH, MEC, and CAMA (n = 8,728)

**Table S4.** Associations of the 180-SNP and 71-SNP PRS with genetic ancestry

| | Indigenous American | | European | | African | |
|---|---|---|---|---|---|---|
| PRS | β (95% CI)* | P-value | β (95% CI)* | P-value | β (95% CI)* | P-value |
| 180-SNP† | -0.002 (-0.006 to 0.002) | 0.40 | 0 (-0.005 to 0.004) | 0.72 | 0.002 (0 to 0.004) | 0.02 |
| 71-SNP‡ | -0.018 (-0.023 to -0.013) | <0.001 | 0.013 (0.008 to 0.019) | <0.001 | 0.005 (0.003 to 0.007) | <0.001 |

Abbreviations: CI = confidence interval; PRS = polygenic risk score

* Beta coefficient and 95% CI from linear regression of PRS (independent variable) on genetic ancestry (dependent variable), adjusted for study. Beta values correspond to change in genetic ancestry on 0 to 1 scale per 1 standard deviation increase in PRS.
† Calculated in controls from 7 datasets, excluding COH/CCGCRN data (n = 7,622)
‡ Calculated in controls from all datasets (n = 7,927)

**Table S5.** Association between 71-SNP PRS and breast cancer risk (COH/CCGCRN participants excluded)

| | Controls | Cases | OR (95% CI)† | P-trend‡ |
|---|---|---|---|---|
| Continuous (per standard deviation) | 7622 | 4658 | 1.49 (1.43 to 1.55) | |
| Percentiles of PRS | | | | <0.001 |
| <10 | 763 | 232 | 0.55 (0.46 to 0.65) | |
| 10-20 | 762 | 292 | 0.69 (0.59 to 0.81) | |
| 20-30 | 761 | 312 | 0.74 (0.63 to 0.86) | |
| 30-40 | 763 | 353 | 0.83 (0.71 to 0.97) | |
| 40-60 | 1525 | 849 | 1 (referent) | |
| 60-70 | 762 | 536 | 1.26 (1.10 to 1.45) | |
| 70-80 | 761 | 562 | 1.33 (1.16 to 1.52) | |
| 80-90 | 763 | 646 | 1.52 (1.33 to 1.74) | |
| >90 | 762 | 876 | 2.06 (1.82 to 2.35) | |

Abbreviations: CI = confidence interval; OR = odds ratio; SD = standard deviation

* Calculated in case-control analysis in 7 datasets, excluding COH/CCGCRN data (n = 12,280)
† Odds ratio from multivariable logistic regression of PRS adjusted for study and genetic ancestry
‡ P-value for test of linear trend between per-decile estimates

**Table S6.** Areas under the receiver operating characteristic curve and odds ratios per standard deviation of the 180-SNP PRS in Hispanics by quartiles of Indigenous American ancestry, stratified by U.S. Latinas and Latin-American women

| | Controls | Cases | AUROC (95% CI)* | *P*-value† | OR (95% CI)‡ | *P*-value§ |
|---|---|---|---|---|---|---|
| U.S. Latinas\|\| | | | | | | |
| All | 5621 | 1696 | 0.63 (0.62 to 0.65) | | 1.62 (1.52 to 1.71) | |
| Quartiles of IA ancestry (%) | | | | 0.06 | | 0.02 |
| Q1, <0.22 | 1493 | 337 | 0.66 (0.63 to 0.69) | | 1.88 (1.65 to 2.13) | |
| Q2, 0.22-0.34 | 1412 | 417 | 0.61 (0.58 to 0.64) | | 1.51 (1.34 to 1.70) | |
| Q3, 0.34-0.45 | 1345 | 484 | 0.63 (0.60 to 0.66) | | 1.57 (1.40 to 1.75) | |
| Q4, >0.45 | 1371 | 458 | 0.62 (0.59 to 0.65) | | 1.53 (1.38 to 1.70) | |
| Latin-American women¶ | | | | | | |
| All | 2001 | 2962 | 0.62 (0.61 to 0.64) | | 1.54 (1.45 to 1.63) | |
| Quartiles of IA ancestry (%) | | | | 0.35 | | 0.49 |
| Q1, <0.43 | 527 | 714 | 0.61 (0.58 to 0.65) | | 1.52 (1.35 to 1.71) | |
| Q2, 0.43-0.54 | 554 | 687 | 0.63 (0.60 to 0.66) | | 1.55 (1.37 to 1.74) | |
| Q3, 0.54-0.71 | 538 | 703 | 0.61 (0.58 to 0.64) | | 1.46 (1.30 to 1.64) | |
| Q4, >0.71 | 382 | 858 | 0.64 (0.60 to 0.67) | | 1.62 (1.43 to 1.82) | |

Abbreviations: AUROC = area under receiver operating characteristic curve; CI = confidence interval; IA = Indigenous American

* AUROC from multivariable logistic regression of PRS adjusted for study and genetic ancestry
† P-value for test of equality of AUROCs between Q1 and Q4 of IA ancestry
‡ Odds ratio per standard deviation increment of PRS adjusted for study and genetic ancestry
§ P-value for comparison of OR of PRS between Q1 and Q4 of IA ancestry using Wald test of linear hypothesis
|| Women from U.S. studies: SFBCS/NC-BCFR, RPGEH, MEC (total n = 7,317)
¶ Women from Latin-American studies: CAMA, Peru, COLUMBUS - Colombia, COLUMBUS – Mexico (total n = 4,963)

**Supplementary Figure S1.** Distribution of log-normalized 180-SNP PRS in 4,658 cases and 7,622 controls

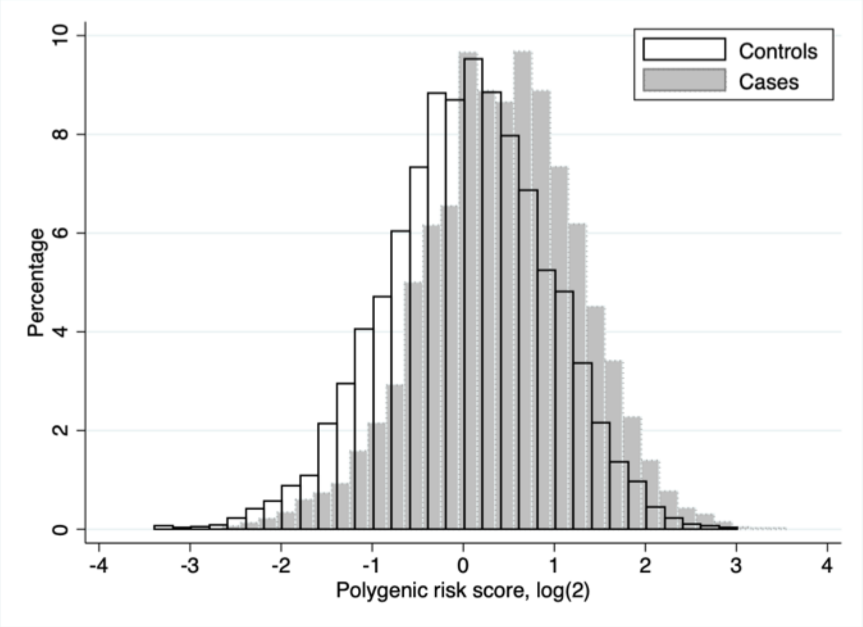

**Supplementary Figure S2.** Ancestry distribution by study and case/control status

A. SFBCS/NC-BCFR

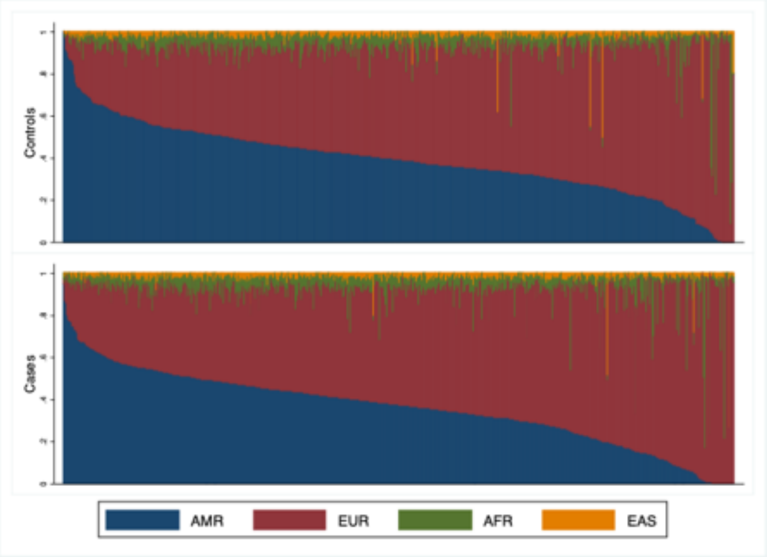

B. RPGEH

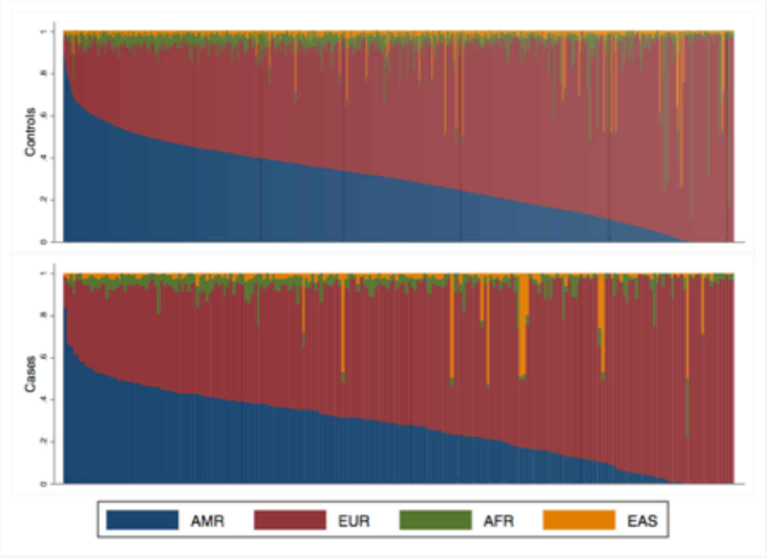

C. MEC

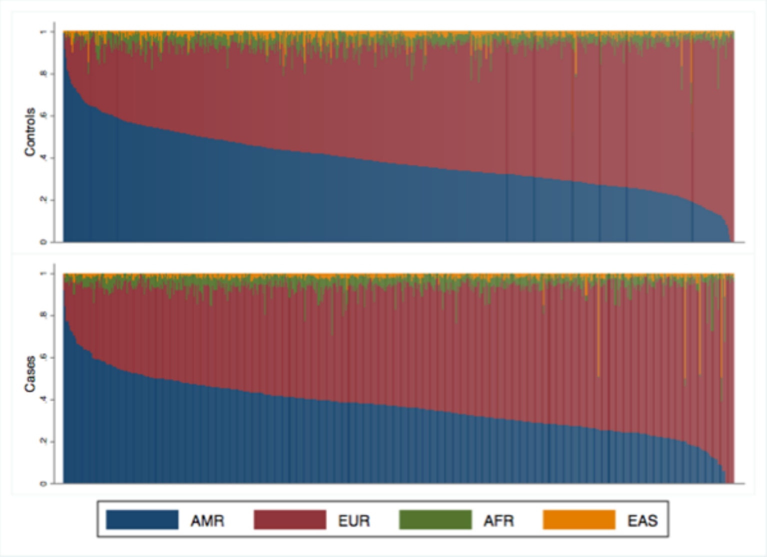

D. CAMA

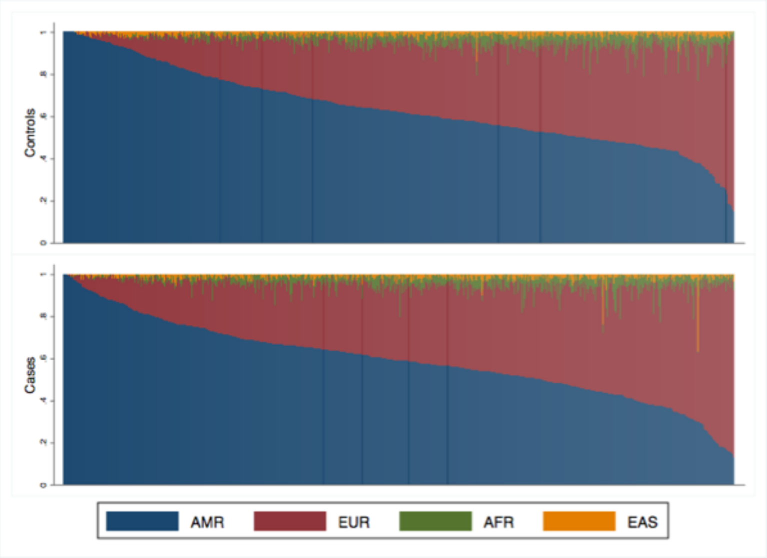

E. COLUMBUS - Colombia

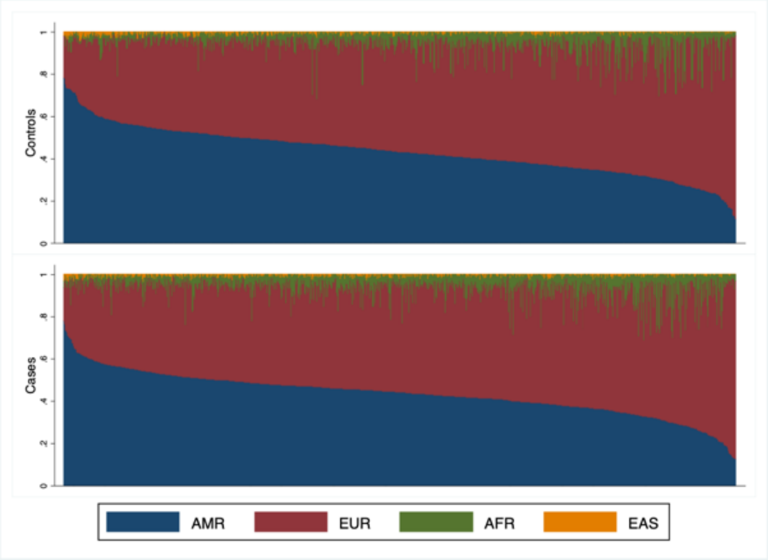

F. COLUMBUS - Mexico

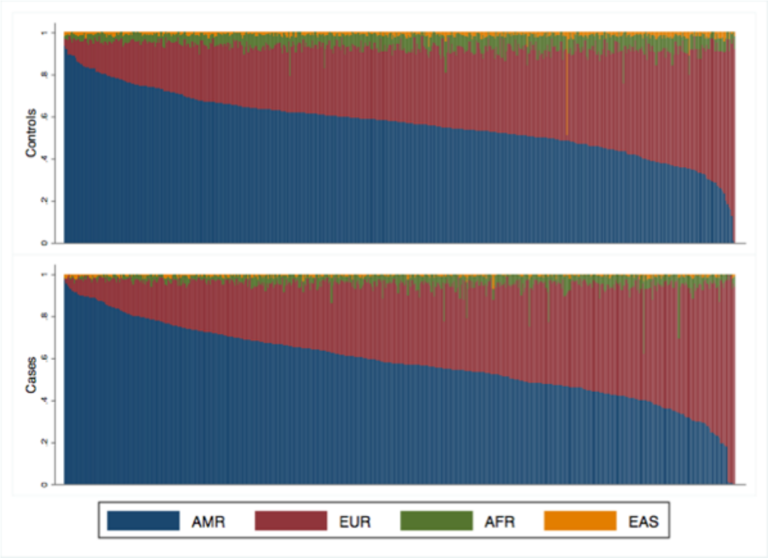

G. Peru

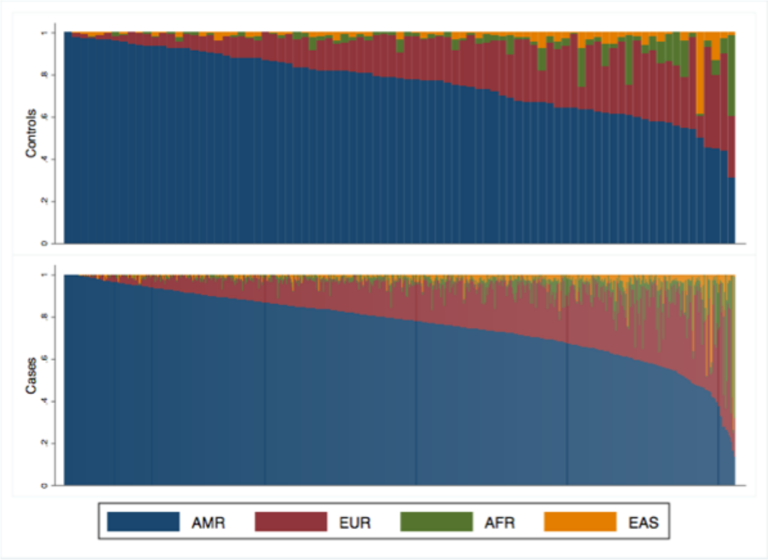

H. CCGRN

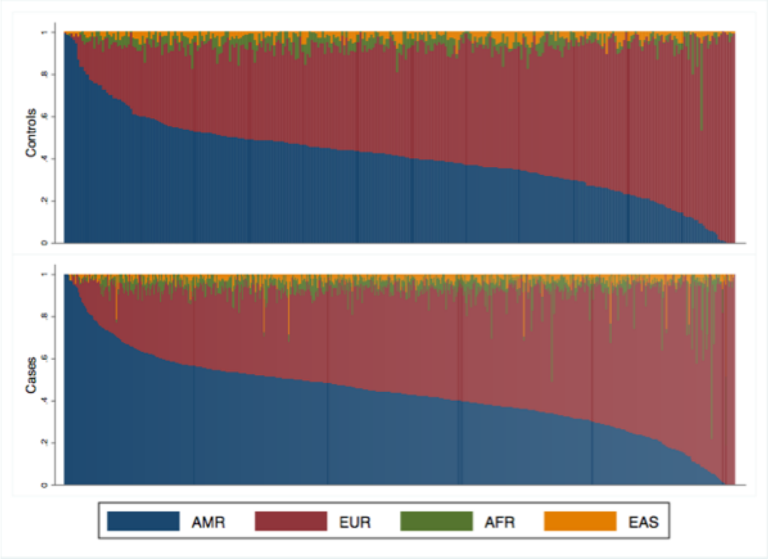
